# Mapping the brain-wide network effects by optogenetic activation of the corpus callosum

**DOI:** 10.1101/810366

**Authors:** Yi Chen, Filip Sobczak, Patricia Pais-Roldán, Cornelius Schwarz, Alan P. Koretsky, Xin Yu

**Author notes:** Corresponding author: Dr. Xin Yu, Address: Max-Planck-Ring 11, 72076, Tuebingen, Germany.

## Abstract

The optogenetically driven manipulation of circuit-specific activity enabled functional causality studies in animals, but its global effect on the brain is rarely reported. Here, we applied simultaneous fMRI with calcium recording to map brain-wide activity by optogenetic activation of fibers running in one orientation along the corpus callosum(CC) connecting the barrel cortex(BC). Robust positive BOLD signals were detected in the ipsilateral BC due to antidromic activity, which spread to ipsilateral motor cortex(MC) and posterior thalamus(PO). In the orthodromic target (contralateral barrel cortex), positive BOLD signals were reliably evoked by 2Hz light pulses, whereas 40Hz light pulses led to a reversed sign of BOLD - indicative of CC-mediated inhibition. This presumed optogenetic CC-mediated inhibition was further elucidated by pairing light with peripheral whisker stimulation at varied inter-stimulus intervals. Whisker induced positive BOLD, and calcium signals were reduced at inter-stimulus intervals of 50/100ms. The calcium-amplitude modulation (AM)-based correlation with whole-brain fMRI signal revealed that the inhibitory effects spread to contralateral BC as well as ipsilateral MC and PO. This work raises the need of fMRI to elucidate the brain-wide network activation in response to projection-specific optogenetic stimulation.

## INTRODUCTION

The genetic expression of channelrhodopsin (ChR2) has been extensively applied to target specific cell types to ensure the activation of neuronal ensembles of interest [1–4]. Optogenetic tools have revolutionized the strategy to perturb or manipulate the behavior of animals [5–8]. To interpret the linkage of the brain function to specific behavioral readout relies on the assumed circuit-specific manipulation through *in vivo* optogenetic activation [9–12]. Optogenetic activation of numerous brain sites and defined neuronal populations in animals has been very successful to modulate behavior. However, there is a lack of systematic mapping of the result of specific modulation on brain-wide network activity, which may relay and affect the proposed link between function and behavior. Progress in this direction depends on the combined application of methods to explore large scale brain dynamics as well [13–16]. One useful method for this purpose is functional magnetic resonance imaging (fMRI), which has been successfully combined with optogenetics [17–22]. We use here a method that adds GCaMP-mediated calcium recordings through an optical fiber for concurrent fMRI and neuronal calcium signal recording [23–27]. This multi-modal cross-scale brain dynamic mapping scheme allows elucidating network activity upon circuit-specific optogenetic activation on the specific target level as well as across large brain regions [19, 23–25, 27–29].

Corpus callosum (CC), the major neural fiber bundles connecting the two hemispheres, plays a critical role to mediate the interhemispheric cortico-cortical connections [30–32]. Despite the highly-correlated structural anomalies of the CC with a wide range of disorders, e.g., schizophrenia [33, 34], autism [35, 36], epilepsy [37, 38] and mental retardation [39, 40], CC-mediated neural mechanisms are primarily studied in loss-of-function models, such as split-brain/callosotomy or partial callosal lesion [31, 41, 42]. To directly investigate the functional roles of callosal projections on regulating the interhemispheric excitatory-inhibitory balance, both *in vitro* and *in vivo* studies have applied micro-stimulation on one hemisphere or the callosal fiber bundles [43–46], or performed bilateral motor or sensory tasks in both human [47–50] and animal models [50–54]. Since the callosal fibers are reciprocally projecting to two hemispheres, bilateral, ortho-vs. antidromically evoked neural activity has been difficult to disentangle. With optogenetic tools, the callosal projection neurons can be specifically (primarily) labeled with ChR2 from one hemisphere, enabling the unidirectional modulation of callosal activity [55–57]. The optogenetically driven callosal activity has been particularly helpful to disentangle interhemispheric inhibitory effects, e.g., in the auditory cortex [58], prefrontal cortex [59] or hindlimb somatosensory cortex [60]. The goal of the present studies was to widen the view beyond of target–specific excitatory-inhibitory regulation by using multi-modal fMRI platform to characterize the global neural network activity upon optogenetic callosal activation.

In the present study, we implemented the multi-modal fMRI platform with optogenetics to map the CC-mediated inhibition on the brain-wide network dynamics in three consecutive steps. First, we identified the antidromic vs. orthodromic effect of CC-specific optogenetic stimulation. Optogenetic stimulation of callosal fibers connecting the barrel cortex (BC) to the other hemisphere, revealed robust antidromic activation in the ipsilateral BC. In the orthodromic direction, both fMRI and neuronal calcium signals in the contralateral BC indicated strong depression of calcium signals with 40Hz light pulses. Second, we specified the temporal characteristics of this presumptive CC-mediated inhibition on the thalamocortical activation to the BC. The optogenetic CC light pulses were paired with the whisker stimulation electrical pulses at varying intervals from 0 ms to 200 ms in a randomized stimulation scheme. Significant inhibitory effects at 50 ms and 100 ms interval were detected by both fMRI and neural calcium recordings of the right BC activated by whisker stimulation, but little difference was observed in the antidromically evoked fMRI signal in the ipsilateral BC. Thirdly, to further examine the brain-wide activity regulation upon paired optogenetic and whisker stimulation, the concurrent evoked-calcium signals in the contralateral BC was real-time detected at varying conditions and correlated with whole-brain fMRI signals. Besides the contralateral BC, the homologous ventral part of the ipsilateral BC, the motor cortex and posterior thalamus (PO) from the same side of the contralateral BC were detected in the correlation maps, showing amplitude modulation by CC-mediated inhibition at varied time intervals. This study not only specifies the optogenetically driven CC-mediated regulation of the local excitation/inhibition balance but also depicts the power of multi-modal fMRI to characterize the brain-wide network activity associated with circuit-specific optogenetic activations *in vivo*. It highlights a vital aspect of the brain-wide activity for circuit-specific causality studies with optogenetic tools.

## RESULTS

### Antidromic activation by callosal optogenetic stimulation

By injecting the AAV-ChR2 viral vectors into the barrel cortex (BC) of rats, ChR2 can be expressed in callosal projection neurons (CPNs), in particular through their axonal fiber bundles projecting to the contralateral BC (Fig. 1a) [56, 61]. Based on our previous work [62], an MRI-guided robotic arm was used to provide high flexibility to insert the optical fiber and sufficient targeting accuracy on the ∼200 µm callosal fiber bundle for multi-modal fMRI. The most salient BOLD fMRI signal evoked by CC optogenetic stimulation was detected at the ipsilateral BC housing the labeled CPN (n = 8 animals, Fig. 1b, c, 5 Hz light pulses). The antidromically evoked hemodynamic responses to 5 Hz stimulation were significantly stronger than the responses to 2 Hz (Fig. 1d). In addition, antidromic BOLD and local field potential (LFP) signal were evoked by systematically varying laser power, light pulse width, frequency and duration of the optogenetic stimulation (Fig. 1e, Fig. S1 and S2). The fMRI analysis revealed widespread brain activation in the ipsilateral hemisphere, which likely originates from antidromic CPN activity spread by multi-synaptic pathways to the motor cortex and posterior thalamus (Fig. 1f). These widespread ipsilateral effects were readily seen with 5Hz stimulation paradigm but could not be evoked using lower stimulus frequencies.

**Fig. 1.**
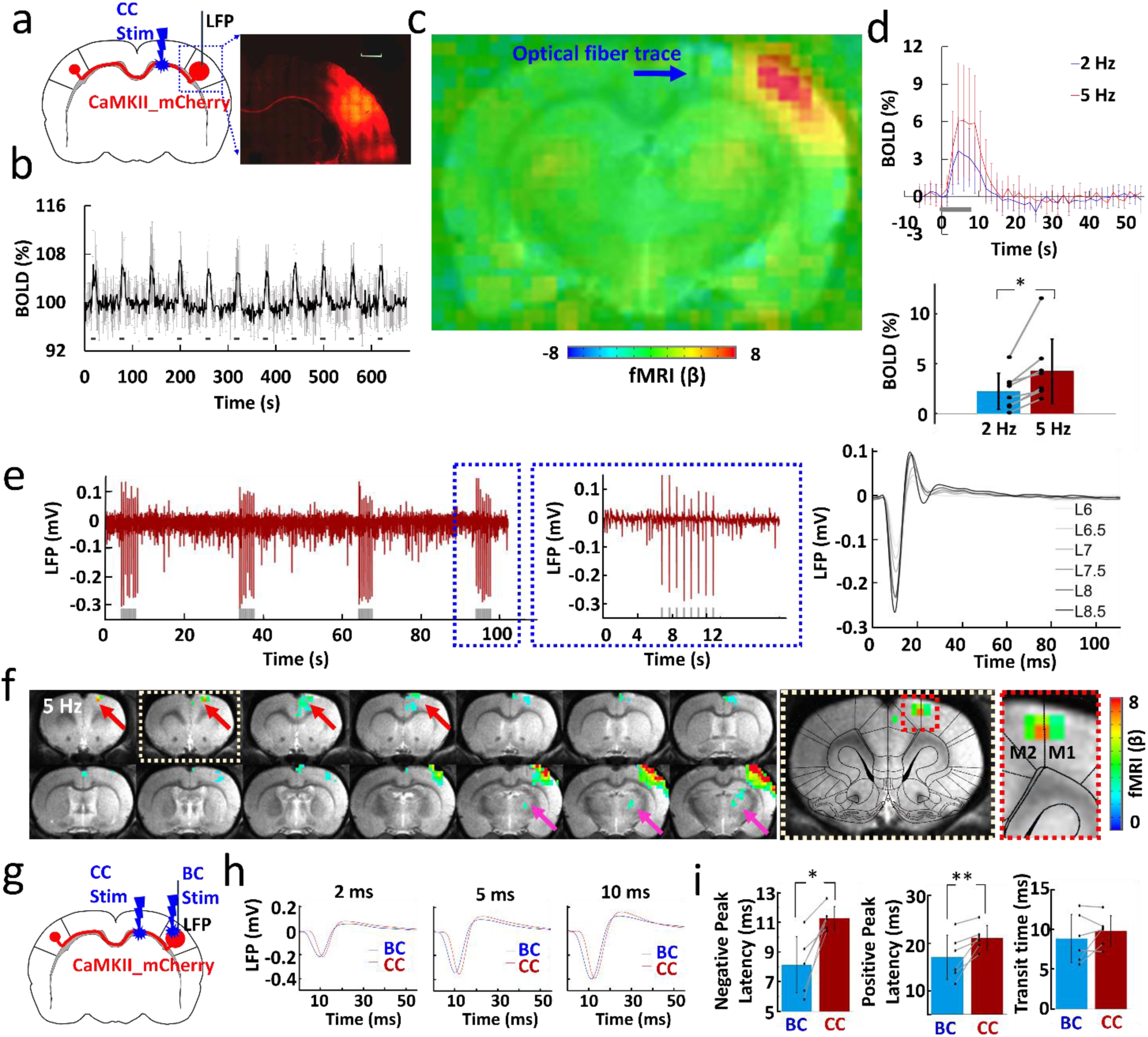
Antidromic activation upon corpus callosum optogenetic stimulation. **a,** *Left:* Schematic of experimental design. *Right:* Representative wide-field fluorescence image illustrating the robust expression of ChR2-mCherry at the injection site (right BC) and along the CC. Medial is to the left. Red, AAV5.CaMKII.ChR2-mCherry. Scale bar, 1 mm. **b,** Average time courses of fMRI signal changes in right BC (n = 8 animals) upon optogenetic stimulation. Error bars represent mean±SD. **c,** Averaged fMRI map showing the strong antidromic activation in BC in the right hemisphere with fiber optic trace (blue arrow) during optogenetic stimulation of CC from 8 rats of block design: 8 s on/ 52 s off, 11 epochs, 10 ms light pulse, 5 Hz. **d,** *Top:* Averaged BOLD signals upon different stimulation frequencies (2 Hz in blue, 5 Hz in red). Error bars represent mean±SD. *Lower:* Mean amplitudes of the BOLD signals (0-10.5 s) for 2 Hz and 5 Hz (n=8, paired t test, *p=0.006). Error bars represent mean±SD. **e**, *Left:* The representative local field potential (LFP) for antidromic activation (gray lines, light pulses). *Right:* Laser power dependent LFPs (pulse width, 10 ms). **f**, Representative BOLD map showing the activity in the projected motor cortex (red arrows) and posterior thalamus (magenta arrow) from the antidromic activity in the BC. Broken boxes showed the enlarged view of projected motor cortex (GLM-based t-statistics in AFNI is used. P (corrected)=0.0319). **g,** Schematic of experimental design. **h,** The representative LFP for direct BC light stimulation (blue) and antidromic activation (red) recorded with light pulse durations of 2, 5 and 10 ms. **i,** The different peak latency and transit time for the LFP induced by CC and BC light stimulation (n = 6 animals, paired t test, *p = 0.002, **p = 0.009, pulse width, 10 ms). Error bars represent mean±SD.

Next, we examined the temporal characteristics of the antidromic activity. In general, CC-mediated antidromic LFP responses in BC to different pulse widths and frequencies were similar to the responses observed when BC was directly activated (Fig. S3 and S4). Likewise, the whole-brain BOLD signal showed time courses and distributions as reported earlier with direct BC stimulation [17, 63]. We were concerned that the stimulation light could have activated CPN directly in the BC. To test this concern, we recorded the LFPs evoked by optogenetic CC and direct BC stimulation in the same rat (Fig. 1g, h), and found that the latency of the response was systematically higher for CC as compared to BC stimulation (negative peak latency: BC: 8.13 ± 1.89 ms, CC: 11.27 ± 0.78 ms; positive peak latency: BC: 17.00 ± 4.65 ms, CC: 21.07 ± 2.60 ms; n = 6 animals, paired t test, *p = 0.002, **p = 0.009) (Fig. 1i). Otherwise the time course of the LFP response was similar, which showed, firstly that CC stimulation is likely stimulating the CC axons as intended, and secondly that BC is activated in a very similar way by CPNs as with direct stimulation. In support of this conclusion, we found that optogenetically activating callosal fibers from the other hemisphere (opposite to the virus injection site) readily showed latency differences (Fig. S5), as expected from the axonal conduction delays of the transmission of the electrical impulses [64, 65].

### Orthodromic activation by callosal optogenetic stimulation

Compared to antidromically evoked activity, the BOLD signal in the contralateral hemisphere evoked by orthodromic stimulation was smaller, and the stimulus-response relationship was different. For instance, quite different from the antidromic situation, the BOLD signal observed with 2 Hz optogenetic CC stimulation was stronger than that with 5 Hz (Fig. S6). To investigate the CC-mediated corticocortical interaction in the contralateral hemisphere, we injected the Syn-GCaMP6f and CaMKII-ChR2-mCherry into the left and right BC, respectively, and recorded both calcium and LFP signal upon optogenetic CC stimulation (Fig. 2a, b). Here we focused on layer 5 (Fig. 2b), because it is the main target lamina of corpus callosum projections [58, 66], as well as the main output layer of the barrel cortex [67]. Fig. 2c shows the frequency-dependent orthodromic calcium signals from one representative rat. As mentioned before evoked calcium transients appeared in strict frequency-dependent fashion. A strong transient was detected following each light pulse at 2 Hz, while at higher frequencies, only the first pulse triggered a full-fledged calcium response (Fig. 2c). The subsequent pulse responses were depressed or missing entirely and gave way to a slow decrement in fluorescence (Fig. 2c). The decrement of Ca^2+^ signal was constantly present throughout the entire stimulus interval (see gray bar 40 Hz stimulation in Fig. 2c), and slowly relaxed back to baseline only after the end of stimulation. Simultaneous LFP and calcium recordings in a representative rat shared the same pattern, strengthening the notion of a strong suppression of responses at higher stimulus frequencies (Fig. S7), and offering an explanation for the likewise decreased orthodromic BOLD signals at 5 Hz (Fig. S6). The calcium baseline drift for 40 Hz was reproduced in animals and was quantified in Fig. 2d, suggesting a highly robust corticocortical inhibition effect as previously reported by electrophysiological recording [59, 68, 69]. The evoked LFP and calcium signals dependencies on the laser power, light pulse width and duration provide strong evidence for reliable detection of the orthodromic activity (Fig. S8-10). It is noteworthy that the CC-mediated orthodromic activity shows different response patterns for both LFP/calcium and fMRI signals from the antidromic activity, indicating a distinct impact on the local excitation-inhibition balance through the CC-mediated inputs.

**Fig. 2.**
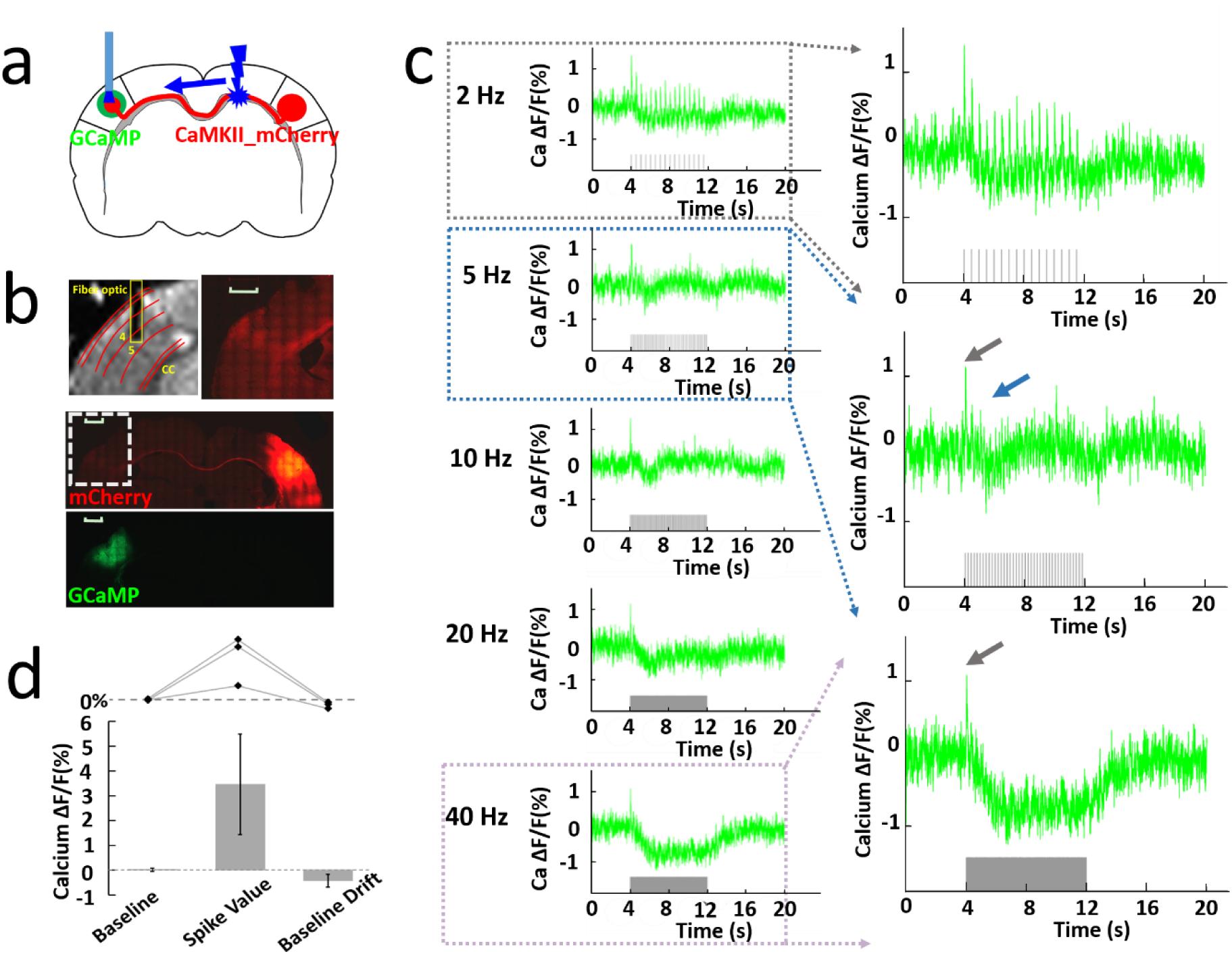
Orthodromic activation upon corpus callosum optogenetic stimulation. **a,** Schematic of experimental design. **b,** *Top left:* Representative RARE anatomical image used to identify the optical fiber location for calcium signal recording in the layer V of barrel cortex. Top right: Enlarged immunostaining image illustrating the ChR2-mCherry expression in the left hemisphere (opposite to the injection site). *Middle:* Representative wide-field fluorescence image illustrating robust ChR2-mCherry at the injection site (right BC) and along the axonal fibers to the other hemisphere. Red, AAV5.CaMKII.ChR2-mCherry. *Bottom:* The immunostaining image illustrating robust GCaMP6f expression (green) in the left barrel cortex. Scale bar, 1 mm. **c,** *Left:* Representative calcium signal changes upon 8 s of orthodromic activation responses to 2, 5, 10, 20 and 40 Hz stimulation. *Right:* Enlarged calcium signal changes responses to 2, 5 and 40 Hz stimulation. **d,** The analysis of calcium baseline, spike value and baseline drift from 3 animals. Error bars represent mean±SD.

### The CC-mediated inhibitory effects on the sensory-evoked cortical activity

Next, we investigated the effect of CC-mediated suppression on sensory-evoked cortical activity. The optogenetic light pulse train (‘O’, 2 Hz, 16 pulses in 8 s) for CC optogenetic stimulation was delivered at time intervals of 0, 50, 100 and 200ms after stimulating the primary afferents in the whisker pad with a microstimulation pulse train (‘W’, 2 Hz, 16 pulses). In total 6 conditions (W, O, OW, O50W, O100W and O200W, OxW means optogenetic pulse leads the whisker stimulation pulse for “x” ms) were delivered in trials of randomized order (Fig. 3a) using the multi-model fMRI platform (Fig. 3b). Typical raw calcium signals and stimulation design are shown in Fig. 3c with a W condition leading the other randomized 12 epochs (6 conditions repeated twice in a randomized order). We found a strong suppression of BOLD in the orthodromic direction with latencies of 50 and 100 ms (Fig. 3d, g). The suppression was partially recovered at the O200W condition. This phenomenon was absent on the antidromic side (Fig, 3g). A similar picture emerged with averaged calcium signals recorded in layer 5 of the contralateral BC (Fig. 3e). Ca^2+^ signals and BOLD were highly correlated (Fig. 3g, e), showing reduced calcium percentage changes at O50W and O100W conditions across animals (Fig. 3f). Normalizing both signals to the whisker-only (W) condition (Fig. S11), we find the mean signal changes of BOLD from 100% (W) to 107.3%, 59.2%, 56.8% and 100.4%, while the calcium signal changed from 100% (W) to 127.8%, 45.2%, 59.5% and 107.1% at conditions of OW, O50W, O100W, and O200W, respectively (Fig. 3h). To investigate the temporal features of the interaction on a more precise scale, we refined the stimulus intervals for whisker stimulation by adding 10 and 25 ms conditions (W, OW, O10W, O25W, O50W, O100W, and O200W) in another group of rats (Fig. 3i and Fig. S12). Again similar patterns emerged as seen before (Fig. 3i and Fig. S12). For O10W, no significant difference was observed in comparison to the OW condition, but the calcium responses at O25W were significantly lower than the OW condition (Fig. S12). As reported from *in vitro* CC electrical stimulation studies by Kawaguchi et al. [44], CC stimulation leads to two inhibitory postsynaptic potential (IPSP) peaks (the earlier peak at ∼30 ms, and the later peak at ∼180 ms), which could underlie the inhibitory effects at O25W and the later recovery at O200W to different extents. Furthermore, the simultaneous LFP and calcium recording confirmed the time-interval specific inhibitory effects by direct optogenetic CC stimulation to modulate the sensory-evoked cortical activity pattern in the BC (Fig. S13). These results are consistent with results using whisker, forepaw, and visual stimulation in rodents and human studies [47–54].

**Fig. 3.**
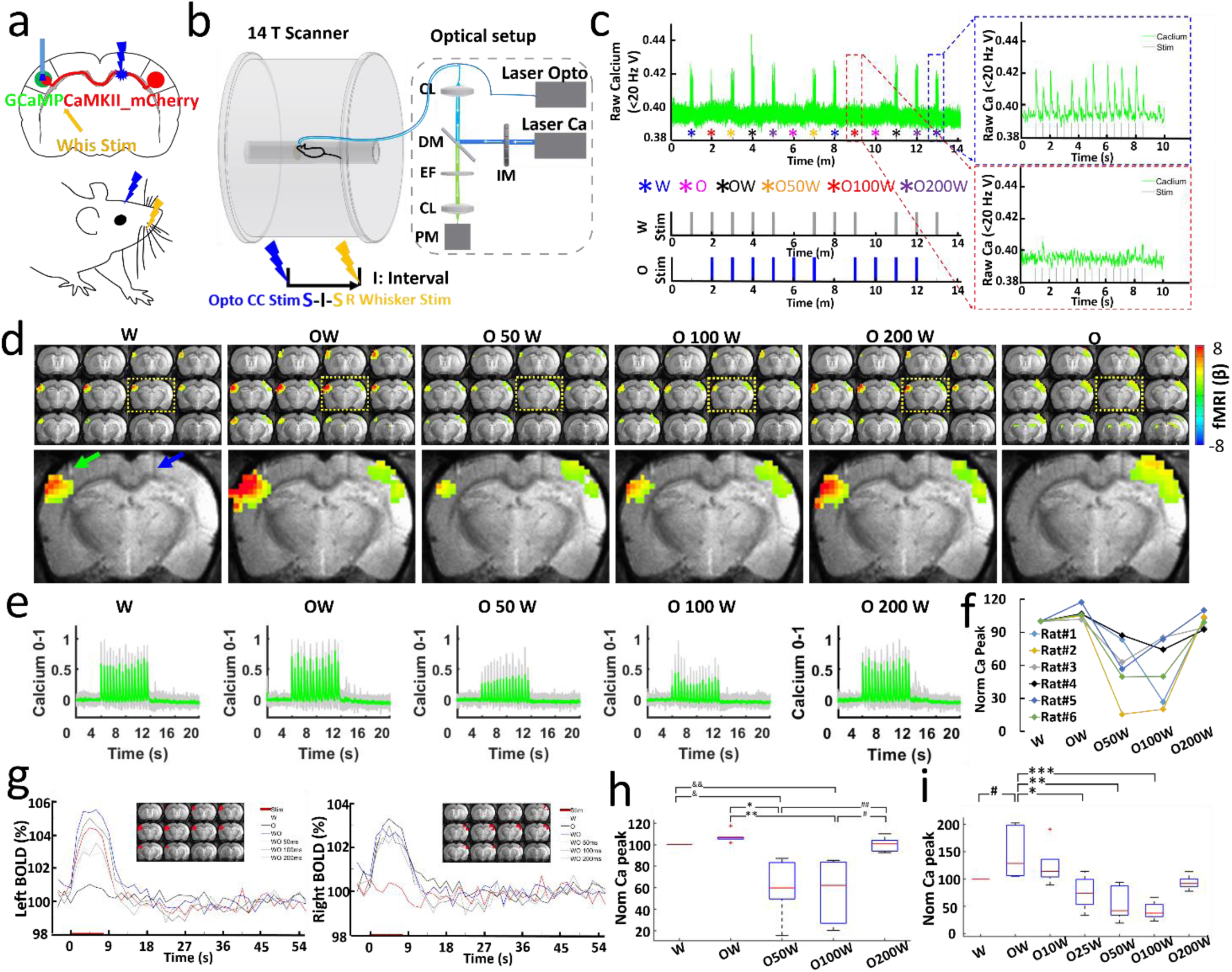
Simultaneous measurement of BOLD and calcium signals during CC optogenetic stimulation and electrical whisker stimulation with varying time intervals. **a,** Stimulation scheme. There are 6 conditions, whisker stimuli only (W), CC stimuli only (O), CC stimuli and whisker stimuli together (OW), CC stimuli and 50 ms, 100ms, 200 ms delayed whisker stimuli (O50W, O100W, O200W). **b,** Schematic drawing of the experimental setup to conduct optogenetic fMRI with simultaneous fiber-optic calcium recording. CL: Coupling Lens, DM: Dichroic Mirror, EF: Emission Filter, PM: Photomultiplier, IM: Intensity Modulation. **c,** Typical calcium signals for condition W (blue dash box) and O100W (red dash box) from a representative rat. **d,** *Top:* Averaged fMRI map of brain-wide activity for 6 conditions across 6 rats (GLM-based t-statistics in AFNI is used. p (corrected) < 0.01) of block design: 8 s on/ 52 s off, 13 epochs, 20 ms light pulse, 2 Hz, 5-39 mw. *Bottom:* Enlarged brain slice showing the differences of BOLD mapping in BC in both hemispheres with fiber optic trace for optogenetic stimulation (blue arrow) and calcium recording fiber (green arrow). **e,** Averaged normalized calcium signal in left BC, grey lines showing the individual normalized calcium signal from 6 rats (Trials # = 29, details see Methods, table 1). **f,** Normalized calcium signal for an individual rat as a function of conditions: W, OW, O50W, O100W, O200W. **g,** *Left:* Averaged BOLD changes in the ROI (red region on anatomical images) in the left BC induced by whisker stimulation. *Right:* averaged BOLD changes in the ROI (red region on anatomical images) in the right BC induced by CC stimulation. **h,** Averaged normalized calcium signal changes across 6 rats modulated by stimulus time intervals (ANOVA, p < 0.01). **i,** Averaged normalized calcium signal changes across 4 rats modulated by stimulus time intervals (ANOVA, p < 0.03).

**Table 1.** The number of trials acquired for 6 conditions.

|  | Rat#1 | Rat #2 | Rat #3 | Rat #4 | Rat #5 | Rat #6 |
| --- | --- | --- | --- | --- | --- | --- |
| **Trials** | 6 | 4 | 4 | 4 | 6 | 5 |
| **Acquiring Time** | 79m 30s | 53m | 53m | 53m | 79m 30s | 66m 15s |

**Table 2.**
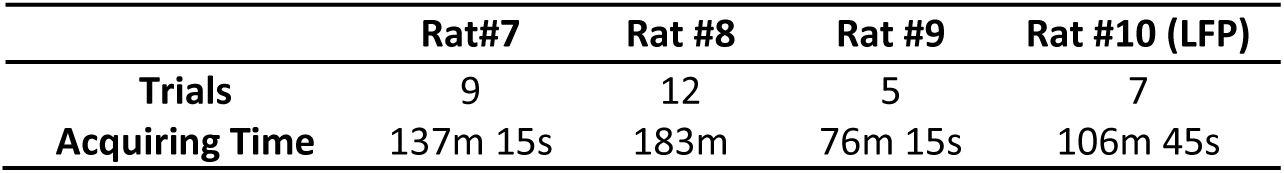
The number of trials acquired with refined stimulus design.

**Table 3.**
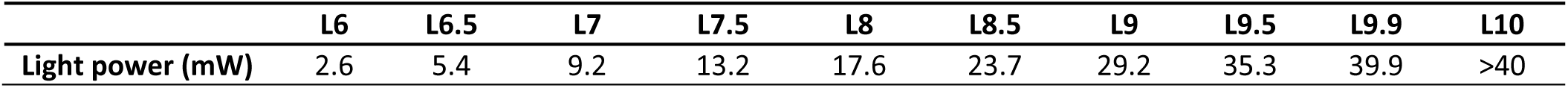
Light power for optogenetic stimulation.

|  | **L6** | **L6.5** | **L7** | **L7.5** | **L8** | **L8.5** | **L9** | **L9.5** | **L9.9** | **L10** |
| --- | --- | --- | --- | --- | --- | --- | --- | --- | --- | --- |
| **Light power (mW)** | 2.6 | 5.4 | 9.2 | 13.2 | 17.6 | 23.7 | 29.2 | 35.3 | 39.9 | >40 |

### Global network mapping based on the optogenetically-driven CC-mediated inhibitory effects

The 3D fMRI data with concurrent calcium signal acquired at different conditions with CC and whisker stimulation allowed analyzing the global effect of the optogenetically-driven CC-mediated inhibition. To this end, the calcium signal amplitude modulation (AM) factor was applied to the ideal function produced by the general linear model (GLM), which was correlated with the 3D fMRI time course (Fig. S14) [24, 70]. As shown in Fig S14, the calcium-AM regressor is derived from the stimulation-driven ideal function, of which the GLM analysis leads to a AM-specific correlation with the whole brain fMRI signal. Thus, the calcium-based AM-correlation with the entire brain generated a map of global brain dynamic changes related to specific CC-mediated inhibition effect. The strongest correlation was found in the left BC (Fig 4). A positive correlation was further observed in the ipsilateral motor cortex and posterior thalamus (PO), which are projection targets of the BC, as well as the vental right BC (Fig. 4a, b, c). We next extracted the time courses from the highlighted ROIs to examine the changes of the fMRI signals at different conditions. The averaged time courses from the right BC ROI reflected the patterns seen in the orthodromically affected BC before. In these conditions (O50W and O100W), the BOLD signals were reduced with respect to the other conditions (Fig. 4d). Similar patterns of BOLD responses were detected in the MC (Fig. 4e) as well as the PO (Fig. 4f) directly connected to the left BC. It is noteworthy that the positively correlated right BC area was not overlapping with cortical areas housing the CPNs (Fig. 3). In summary, these results demonstrate that the global network is modulated with the CC-specific evoked activity in BC. The specificity of CPN precludes the possibility that MC and PO might have integrated the callosal and sensory input independently of BC.

**Fig. 4.**
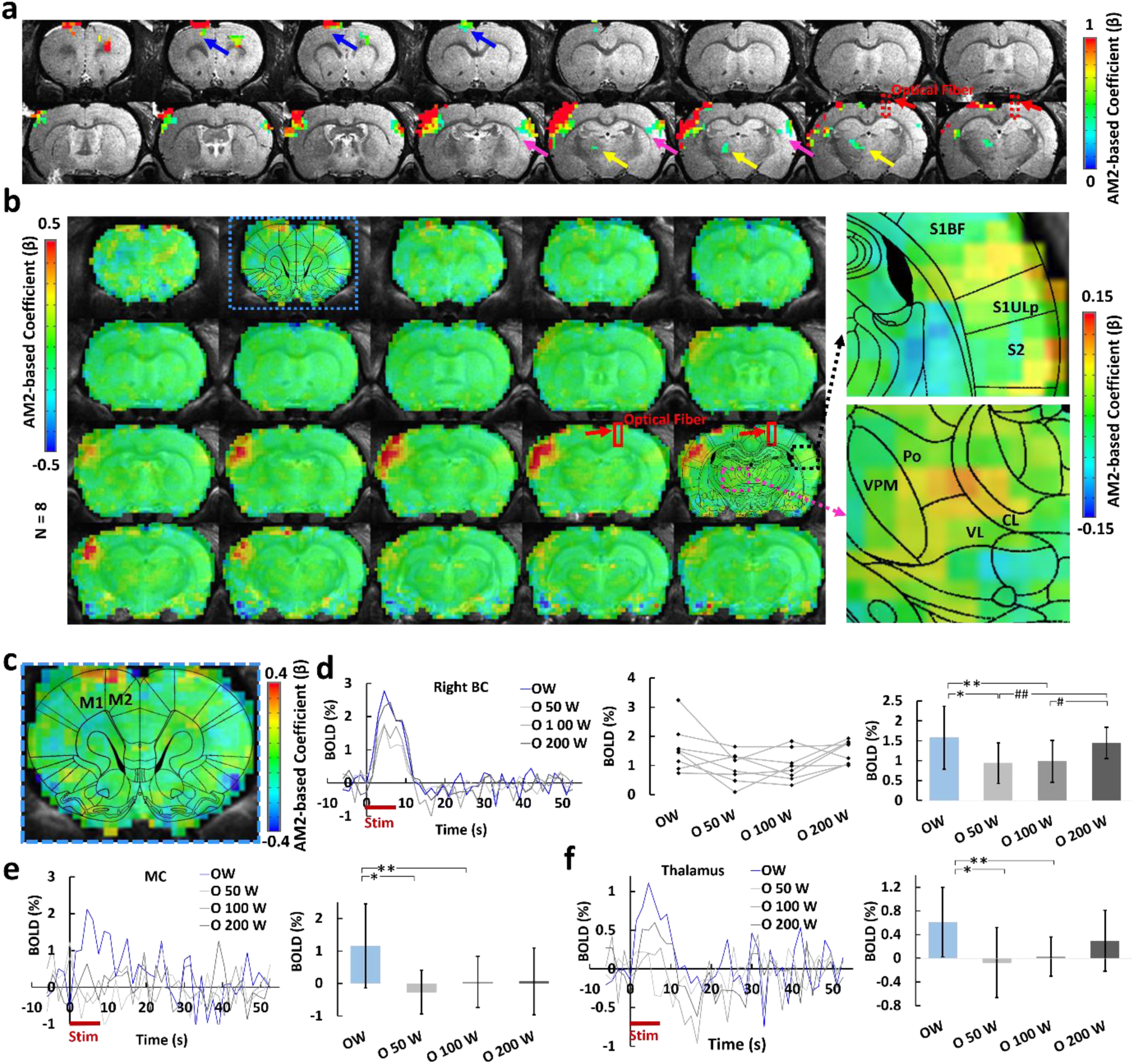
Calcium AM-based whole brain BOLD correlation analysis. **a**, one representative animal map overlaid on the anatomical image with a statistical threshold (p (corrected) < 0.05, cluster size > 15 voxels, MC, blue arrows, BC, magenta arrows, PO, yellow arros, optical fiber trace, red arrows). **b**, Group-averaged correlation maps show the spatial distribution of the positive correlation located at left BC, MC, as well as the PO by overlying with the brain atlas (red square, optical fiber traces, right panel: the enlarged images of the correlation map overlaid on the brain atlas). **c**, Enlarged correlation map shows the positive correlation at the MC. **d**, *Left*: Averaged time courses from the right BC at different conditions (n = 8 rats). *middle*: Mean amplitudes of the BOLD signals (0-10.5 s) for individual rats. *Right*: Averaged amplitudes of the BOLD signals (0-10.5 s, mean±SD, ANOVA, *p = 0.027, **p = 0.004, #p = 0.030, ##p = 0.003). **e**, *Left:* averaged time courses from the MC (n = 8 rats). *Right:* Averaged amplitudes of the BOLD signals (0-10.5 s, mean±SD, ANOVA, *p = 0.005, **p = 0.01). **f**, *Left:* Averaged time courses from the PO (n = 8 rats). *Right*: Averaged amplitudes of the BOLD signal (0-10.5 s, mean±SD, ANOVA, *p = 0.009, **p = 0.012). W: whisker stimulation only, OW: simultaneous optical and whisker stimulation, O[x]W optical stimulation followed by [x] ms-delayed whisker stimulation.

## DISCUSSION

We have performed simultaneous BOLD-fMRI and calcium recording in combination with callosal-circuit specific optogenetic stimulation to map the brain-wide network activation. The robust BOLD signal due to the antidromic activity was detected in the ipsilateral BC, which also led to fMRI detection in the ipsilateral MC and PO region with the higher frequency stimulus. In contrast, the positive BOLD signal through the CC-orthodromic activity was only reliably observed at the lower frequency optogenetic stimulus. With the 40Hz light pulses, the calcium baseline suppression was detected and interpreted to be due to the CC-mediated cortico-cortical inhibitory effect. To further test this CC-mediated inhibition was further paired with the whisker stimulation paradigm at varying inter-stimulus intervals from 0 ms to 200 ms, showing significant suppression at the O50W and O100W conditions in the left BC by the concurrent fMRI and calcium recording. By extracting the event-dependent calcium peak amplitudes at varied conditions as a regressor, an amplitude modulation (AM)-based correlation map revealed the brain-wide inhibitory effects spreading through the ventral border of the right BC and the left MC and PO. Thus, the multi-modal fMRI platform provides a thorough brain-wide network activation maps for the CC-specific optogenetic stimulation.

The observation of strong antidromic propagation by callosal optogenetic stimulation and related synaptic spread of activity presents a caveat for the conclusion of circuit specificity for *in vivo* optogenetic studies. In particular, when neuronal projection terminals labeled with ChR-2 from neurons located at specific functional nuclei are targeted, possible spreading network activity from the antidromically activated brain sites need to be considered. In our experiments, BOLD signals were detected in both MC and PO projected from the antidromically activated BC (at 5Hz light pulses), indicating a (for the experimental purpose unintended) wide-spread optogenetic activation pattern in the brain-wide network (Fig 1e). This spread is likely due to synaptic propagation via activated local or regional axon collaterals of CPNs [71–75]. For the present spread into motor and sensorimotor structures, deep layer CPN with long-range projections into sensorimotor brain areas are likely involved [76]. In addition, multi-synaptic pathways, involving either cortico-cortical or cortico-thalamic projections may have contributed to the spread brain-wide activation [77, 78]. In conclusion, it is mandatory to consider brain-wide activation patterns, even in case of application of highly circuit-specific optogenetic activation schemes.

Certainly, the optogenetic callosal fiber activation also elicits the specific unidirectional callosal orthodromic activity as well, similar to earlier reports[43, 44, 46]. In addition, the optogenetic activation of the callosal projection terminals from brain slices leads to better characterization of the excitatory and inhibitory circuit regulation by callosal inputs [56, 58–60, 79]. Our observations further support the non-linear neurovascular coupling events with the optical intrinsic signal measurements and laser-doppler flowmetry upon the optogenetic or electrical CC stimulation [43, 55]. In our study, the fact that orthodromic BOLD signals were readily observed with low-frequency stimulation (2 Hz), but were strongly reduced at the higher frequency (5Hz), reveals a critical non-linear manner of the hemodynamic responses driven the the CC-mediated neuronal activation(Fig. 1f, 2c, and Fig. S6). We show here that peripheral whisker stimulation is well suited to study the suppressive effects of orthodromically conveyed activity specific to the callous, which is not possible using *in vivo* bilateral stimulation paradigms in rodents [50–54] or bilateral motor or visual tasks in humans [47–49] where other pathways maybe involved. In particular, CC-induced orthodromic activity of L5 pyramidal neurons evoked a calcium transient followed by marked depression of calcium signals responding to light pulses on CC (Fig. 2c,d) (consistent with the optogenetic results in brain slices [59]). Electrophysiology in brain slices has elucidated that CC-mediated glutamatergic excitatory postsynaptic potentials are followed by early GABA_A_- and late GABA_B_-mediated inhibitory postsynaptic potentials lasting for several hundred milliseconds [44–46, 60], strongly suggesting that the depression seen here is partly due to synaptic inhibition. Also, while pairing with 2 Hz whisker stimulation, a time course of the depressive effect around 50-100 ms interval fit the previous finding that local intracortical activation is characterized by activation of long-lasting synaptic GABAergic inhibition [57, 68, 69, 80–82]. In particular, besides the robust inhibition detected in the paired O50W and O100W conditions, a refined temporal scale at the O25W condition further demonstrates the CC-mediated inhibitory effect (Fig. S12), which can be potentially caused by the GABA_A_-mediated early IPSP peak elicited by the direct electrical CC stimulation [44]. The fact that antidromic activity is not susceptible for the paired optogenetic and whisker stimulation (surely due to weaker ipsilateral whisker-evoked activity, but also likely due to the relative strength of antidromic activation), supports the notion that the depression of whisker-evoked activity is due mainly to local (contralateral) interaction of CC-evoked and whisker-evoked activity, rather than to possible CC activity evoked by indirect activation of additional CPNs via antidromic activation.

The whole-brain fMRI with concurrent calcium recording allows accessing brain-wide network effects of CC-mediated inhibition (Fig. 4a, b). In particular, the applicaton of the AM-based GLM allows separating the stimulus-driven reponses from the AM factor, which creates specific correlation maps to the CC-mediated inhibitory effects. The calcium amplitude-modulation (AM)-based correlation map highlighted three brain regions: the ventral part of right BC, the left MC, and PO. The ventral right BC was likely activated by reciprocal callosal connections, the majority of which, as argued above, may have been quenched by the strong antidromic effect via labeled CPNs. In the injection experiments, however, the ventral BC was regularly spared and did not receive virus, and therefore may have been less affected by overriding antidromic activity. Outside BC on the orthodromic side the AM-dependent correlation was detected as well in the right MC and PO. The CC-mediated inhibitory effect on the spatially distinct MC could be caused by the long-range S1-MC projection for sensorimotor integration [78, 83–86]. The direct BOLD activation in the MC was detected by whisker stimulation through the sensorimotor connection [87], which was also shown in the antidromic activity-based spreading activation patterns (Fig 1e). The CC-mediated inhibitory effect on the PO is likely via corticothalamic projections originating from BC layer 5b neurons [88–91]. This finding points at a potential participation of the callosal inputs in the regulation of a wider network of a reciprocal thalamocortical network which mediates BC signals from the other hemisphere for whisking related processing [77, 83, 89, 92–95]. Therefore, besides the antidromically evoked network activation pattern, the orthodromic CC-mediated inhibition generates a brain-wide activity pattern of its own.

In summary, by taking advantage of optogenetics to activate unidirectional callosal fiber, calcium indicators (GCaMP6f) to track specific L5 pyramidal neuronal activity, and simultaneous whole-brain fMRI mapping, this work bridges the scales from the cellular to the whole brain network level for CC-mediated activity. We present a multi-modal fMRI platform to map and analyze the CC-regulated excitation/inhibition balance across multiple scales, which should be useful to decipher brain network dysfunction induced from CC abnormalities. Brain-wide network activation from callosal-circuit optogenetic stimulation underscores the caution to interpret circuit-specific regulatory mechanisms underlying behavioral or functional outcomes with optogenetics in animals.

## Materials and methods

### Animal procedures

The study was performed in accordance with the German Animal Welfare Act (TierSchG) and Animal Welfare Laboratory Animal Ordinance (TierSchVersV). This is in full compliance with the guidelines of the EU Directive on the protection of animals used for scientific purposes (2010/63/EU). The study was reviewed by the ethics commission (§15 TierSchG) and approved by the state authority (Regierungspräsidium, Tübingen, Baden-Württemberg, Germany). A 12-12 hour on/off lighting cycle was maintained to assure undisturbed circadian rhythm. The food and water were obtainable ad libitum. A total of 24 (17 for fMRI and 7 for electrophysiology) male Sprague–Dawley rats were used in this study.

### Viral injection

Intracerebral viral injection was performed in 4-week-old rats to express the viral vectors containing the light-sensitive protein channelrhodopsin-2 (ChR2, for optogenetics) and/or the calcium-sensitive protein (GCaMP, for calcium recording) in neurons. The construct AAV5.Syn.GCaMP6f.WPRE.SV40 was used to express GCaMP in the left BC and the constructs AAV5.CaMKII.hChR2(H134R)-mCherry.WPRE.hGH was used to express ChR2 in the right BC. The stereotaxic coordinates of the injections were ±2.5 mm posterior to Bregma, 5.0 mm lateral to the midline, 0.8-1.4 mm below the cortical surface. Rats were anesthetized with 1.5-2% isoflurane via nose cone and placed on a stereotaxic frame, an incision was made on the scalp and the skull was exposed. Craniotomies were performed with a pneumatic drill so as to cause minimal damage to cortical tissue. A volume of 0.6-0.9 µL and 0.6 µL, for optogenetics and calcium signal recording, respectively, was injected using a 10 µL syringe and 33-gauge needle. The injection rate was controlled by an infusion pump (Pump 11 Elite, Harvard Apparatus, USA). After injection, the needle was left in place for approximately 5 min before being slowly withdrawn. The craniotomies were sealed with bone wax and the skin around the wound was sutured. Rats were subcutaneously injected with antibiotic and painkiller for 3 consecutive days to prevent bacterial infections and relieve postoperative pain.

### Immunohistochemistry

To verify the phenotype of the transfected cells, opsin localization and optical fiber placement, perfused rat brains were fixed overnight in 4% paraformaldehyde and then equilibrated in 15% and 30% sucrose in 0.1 M PBS at 4°C. 30 µm-thick coronal sections were cut on a cryotome (CM3050S, Leica, Germany). Free-floating sections were washed in PBS, mounted on microscope slides, and incubated with DAPI (VectaShield, Vector Laboratories, USA) for 30 mins at room temperature. Wide-field fluorescent images were acquired using a microscope (Zeiss, Germany) for assessment of GCaMP and ChR2 expression in BC. Digital images were minimally processed using ImageJ to enhance brightness and contrast for visualization purposes.

### Optical setup for calcium recordings

A laser was used as excitation light source (OBIS 488LS, Coherent, Germany) with a heat sink to enable laser operation throughout the entire specified temperature range from 10°C to 40°C. The light passed through a continuously variable neutral density filter (NDC-50C-2M-B, Thorlabs, Germany) and was reflected on a dichroic beam splitter (F48-487, AHF analysentechnik AG, Germany). The beam was collected into an AR coated achromatic lens (AC254-030-A, Thorlabs, Germany) fixed on a threaded flexure stage (HCS013, Thorlabs, Germany) mounted on an extension platform (AMA009/M, Thorlabs, Germany) of a fiber launch system (MAX350D/M, Thorlabs, Germany). The laser beam was projected into the fiber and propagated to its tip. The fluorescence emitted by neurons was collected through the fiber tip, propagated back and collimated by the achromatic lens, passed through the dichroic beam splitter and filtered by a band-pass filter (ET525/50M, Chroma, USA) and focused by an AR coated achromatic lens (AC254-030-A, Thorlabs, Germany). A silicon photomultiplier module (MiniSM 10035, SensL, Germany) was applied to detect the emitted fluorescence. The entire optical system was enclosed in a light isolator box. The photomultiplier output was amplified (gain = 100) by a voltage amplifier (DLPVA-100-BLN-S, Femto, Germany), digitized and detected by BIOPAC system (MP150 System, BIOPAC Systems, USA).

### Animal preparation and fiber optic implantation for fMRI

Anesthesia was first induced in the animal with 5% isoflurane in the chamber. The anesthetized rat was intubated using a tracheal tube and a mechanical ventilator (SAR-830, CWE, USA) was used to ventilate animals throughout the whole experiment. Femoral arterial and venous catheterization was performed with polyethylene tubing for blood sampling, drug administration, and constant blood pressure measurements. After the surgery, isoflurane was switched off, and a bolus of the anesthetic alpha-chloralose (80 mg/kg) was infused intravenously. A mixture of alpha-chloralose (26.5 mg/kg/h) and pancuronium (2 mg/kg/h) was constantly infused to maintain the anesthesia/keep the animal anesthetized and reduce motion artifacts.

Before transferring the animal to the MRI scanner, two craniotomies were performed: one for fixed fiber implantation to record calcium signals from BC, and the other one for dynamic insertion of the optical fiber to stimulate the CC using optogenetics (dynamic insertion was achieved by using a remote positioning tool [62]). The animal was placed on a stereotaxic frame, the scalp was opened and two ∼1.5 mm diameter burr holes were drilled on the skull. The dura was carefully removed and an optical fiber with 200 µm core diameter (FT200EMT, Thorlabs, Germany) was inserted into the BC, at coordinates: 2.75-3.3 mm posterior to Bregma, 5.0 mm lateral to the midline, 1.2-1.4 mm below the cortical surface. An adhesive gel was used to secure the calcium recording fiber to the skull. The craniotomy for optogenetics on CC in the other hemisphere, at coordinates: 2.75-3.3 mm posterior to Bregma, 1.8-2.4 mm lateral to the midline, was covered by agarose gel for the robotic arm-driven fiber insertion inside the MRI scanner. The eyes of the rats were covered to prevent stimulation of the visual system during the optogenetic fMRI, which can occur in cases with imperfect coverage or under the strong power of light pulses through tissue.

### Functional MRI acquisition

All images were acquired with a 14.1 T/26 cm horizontal bore magnet interfaced to an Avance III console and equipped with a 12 cm gradient set capable of providing 100 G/cm over a time of 150 µs. A transceiver single-loop surface coil with an inner diameter of 22 mm was placed directly over the rat head to acquire anatomical and fMRI images. Magnetic field homogeneity was optimized first by global shimming for anatomical images and followed by FASTMAP shimming protocol for the EPI sequence. Functional images were acquired with a 3D gradient-echo EPI sequence with the following parameters: Echo Time 11.5 ms, repetition time 1.5 s, FOV 1.92 cm × 1.92 cm × 1.92 cm, matrix size 48 × 48 × 48, spatial resolution 0.4 mm × 0.4 mm × 0.4 mm.

For anatomical reference, the RARE sequence was applied to acquire 48 coronal slices with the same geometry as that of the fMRI images. The paradigm for each trial consisted of 360 dummy scans to reach steady-state, 10 pre-stimulation scans, 5 scans during stimulation (stimulation period 8 s), 35 post-stimulation scans with total 13 epochs and 15 epochs for refined stimulus design (See **Stimulation protocols**).

For fMRI and electrophysiology studies, needle electrodes were placed on whisker pads of the rats, and electric pulses (333 µs duration at 1.5 mA repeated at 3 Hz for 4 seconds) were first used as stimulation to serve as a positive control for the evoked BOLD signal or local field potential/calcium signal. Once that reliable fMRI signals and calcium signals were observed in response to electrical stimulation, optical stimulation was performed. For optogenetic stimulation, square pulses of blue light (473 nm) were delivered using a laser (MBL-III, CNI, China) connected to the 200 µm core optical fiber (FT200EMT, Thorlabs, Germany) and controlled by Master 9 (Master-9, A.M.P.I., Israel) to deliver blue light pulses at 1-40 Hz, 1-20 ms pulse width with 2-8 s duration. The light intensity was tested before each experiment and was calibrated with a power meter (PM20A, Thorlabs, Germany) to emit 0.6 mW to 40 mW from the tip of the optical fiber for CC activation.

### Stimulation protocols

A 2 Hz, 8 s optogenetic stimulus train (O train; 16 pulses to the corpus callosum) was delivered preceding a conditioning stimulus train (W train; same pulse parameters were used, 0.75-1.5 mA) while varying the time interval between stimuli (0, 10, 25, 50, 100 and 200 ms), or without a W train, in a single trial. These stimulation conditions were automatically executed using a laser (MBL-III, CNI, China) and a stimulator (A365 Stimulus Isolator, WPI, USA) triggered by a combination program provided by pulse generator (Master-9, A.M.P.I., Israel), which were precisely synchronized with the start time of the image acquisition sequence in each trial. Each trial consisted of the first fixed whisker stimuli block (W) and 12 blocks randomized for 6 different conditions, W, O, WO, W50O, W100O, W200O, in total 13 min and 15 s for each trial. For refined inter-stimulus intervals design, first fixed whisker stimuli block (W) and 14 blocks randomized for 7 different conditions were used: W, OW, O10W, O25W, O50W, O100W, and O200W, in total 15 min 15 s for each trial. The tables below show the number of continuous trials acquired in this study, as well as light power for optogenetic stimulation.

### Simultaneous calcium recording with electrophysiology

The anesthetic and surgical preparation procedures were similar to the fMRI experiments. For antidromic activity recording experiments in Fig. 1 and Fig. S2-5, tungsten microelectrode (UEWSDDSMCN1M, FHC, USA) was implanted in the right BC to record the LFP from the callosal projection neurons. For orthodromic activity in Fig. 2 and Fig. S7-10, the same kind of tungsten microelectrode was attached to the fiber optic closely, implanted in the left BC, then secured to the skull by an adhesive gel. To calculate the coordinates of optical fiber implantation for CC activation, a FLASH anatomical MRI image was acquired to confirm the virus injection one day before the experiment. The LFP was recorded and amplified through the EEG module of the BIOPAC system (gain factor, 5000, band-pass filter, 0.02-100 Hz, sampling rate, 5,000/s). In parallel, the GCaMP6f-mediated fluorescent signal and blood pressure were digitized and recorded with BIOPAC (MP150 System, BIOPAC Systems, USA) at a sampling rate of 5 kHz. The experiment design and equipment used afterward were similar to the fMRI experiments.

### Data analysis

Acquired data were analyzed using Functional NeuroImages software (AFNI, NIH, USA) and custom-written Matlab (MATLAB, MathWorks, USA) programs for calcium signals. The fiber optical neuronal calcium signals were low-pass filtered at 100 Hz using zero-phase shift digital filtering (filtfilt function in MATLAB). The relative percentage change of fluorescence (ΔF/F) was defined as (F-F_0_)/F_0_, where F_0_ is the baseline, i.e., the average fluorescent signal in a 2 s pre-stimulation window. For Fig. 2d, the spike value is defined by the maximal value for the difference in ΔF/F in a time window 0.3 s after the stimulus, as shown from 40 Hz in Fig. 2c, while the baseline drift is the average calcium signal from 0.3–8 s after the spike recovered to baseline for 40 Hz stimulation. For Fig. 3e, the first epoch for each trial (fixed W condition) was excluded in the data analysis and the calcium signal was averaged for each condition from all the acquired trials for each animal. Each condition was then normalized by the maximum positive deflection of calcium signal alone conditions. For Fig. 3f, h, i, the amplitude peak of the neuronal fluorescent signal in response to 8 s whisker stimulus was calculated as the maximal difference in ΔF/F in a time window 300 ms after stimulus, then normalized to the whisker only (W) condition (100%). The unnormalized amplitude for the difference in ΔF/F for each epoch was used to generate the calcium signal-based regressor (Fig. S14) for fMRI correlation map in Fig. 4.

For evoked fMRI analysis, EPI images were first aligned to anatomical images acquired in the same orientation with the same geometry. The anatomical MRI images were registered to a template across animals, as well as EPI datasets. The baseline signal of EPI images was normalized to 100 for statistical analysis of the multiple trials of EPI time courses. The time courses of the BOLD signal were extracted from regions of interest, e.g., barrel cortex, motor cortex, and posterior thalamus, which were segmented on the anatomical images based on the brain atlas and activation or correlation values. The BOLD amplitude for each condition was defined as the average value for the volumes within the 0-10.5 s following the onset of stimulation (when stimulation duration was 8 s). The hemodynamic response function (HRF) used was the default of the block function of the linear program 3dDeconvolve in AFNI. BLOCK (L, 1) computes a convolution of a square wave of duration L and makes a peak amplitude of block response = 1, with *g*(*t*) = *t*^4^*e*^−4^/[4^4^*e*^−4^] (peak value=1). In this case, each beta weight represents the peak height of the corresponding BLOCK curve for that class, i.e. the beta weight is the magnitude of the response to the entire stimulus block, as shown in Fig. 1, 3 and Fig. S1. The HRF model is defined as follows:

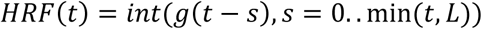

For correlation analysis, a calcium signal amplitude modulated regressor (AM2) based AFNI BLOCK (L, 1) function was used (Fig. S14). The regressor for amplitude modulated response model is as follows:

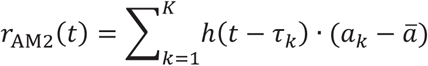

Where *a_k_* = value of *k^th^* auxiliary behavioral information value (ABI), i.e., calcium amplitude value for the difference in ΔF/F for each epoch, and *ā* is the average calcium amplitude value for all the epochs for the individual animal. The statistics and *β* for AM2 regressor make activation map of voxels whole BOLD response vary proportionally to ABI, i.e., the changes in calcium signals for each epoch.

## Author Contributions

X.Y. designed and supervised the research, Y.C. and X.Y. performed animal experiments, Y.C. acquired data, Y.C. analyzed data, A.K., C.S., F.S. and P.P-R. provided conceptual and technical support, X.Y., Y.C., A.K. and C.S. wrote the manuscript.

## Data availability

Excel files are included for each quantitative plot included in the main figures. All other data generated during this study are available from the corresponding author upon reasonable request.

## Code availability

The Analysis of Functional NeuroImages software (AFNI, NIH, USA) and Matlab (MATLAB, MathWorks, USA) were used to process the fMRI and simultaneously acquired calcium signals, respectively. The relevant source codes can be downloaded through https://afni.nimh.nih.gov/afni/. The related image processing codes are available from the corresponding author upon reasonable request.

## ACKNOWLEDGEMENTS

The financial support of the NIH grant (1RF1NS113278), Max-Planck-Society, DFG (YU 215/3-1)), BMBF (01GQ1702) and the China Scholarship Council (Ph.D. fellowship to Y. Chen) are gratefully acknowledged. We thank Dr. N. Avdievitch and Ms. H. Schulz for technical support, Dr. P. Douay, Mrs. R. König, Dr. E. Weiler, Ms. S. Fischer and Mrs. M. Pitscheider for animal support, the AFNI team for the software support, the Genetically-Encoded Neuronal Indicator and Effector Program and the Janelia Farm Research Campus for kindly providing viral plasmids.

---

**Supplementary Figure 1.**
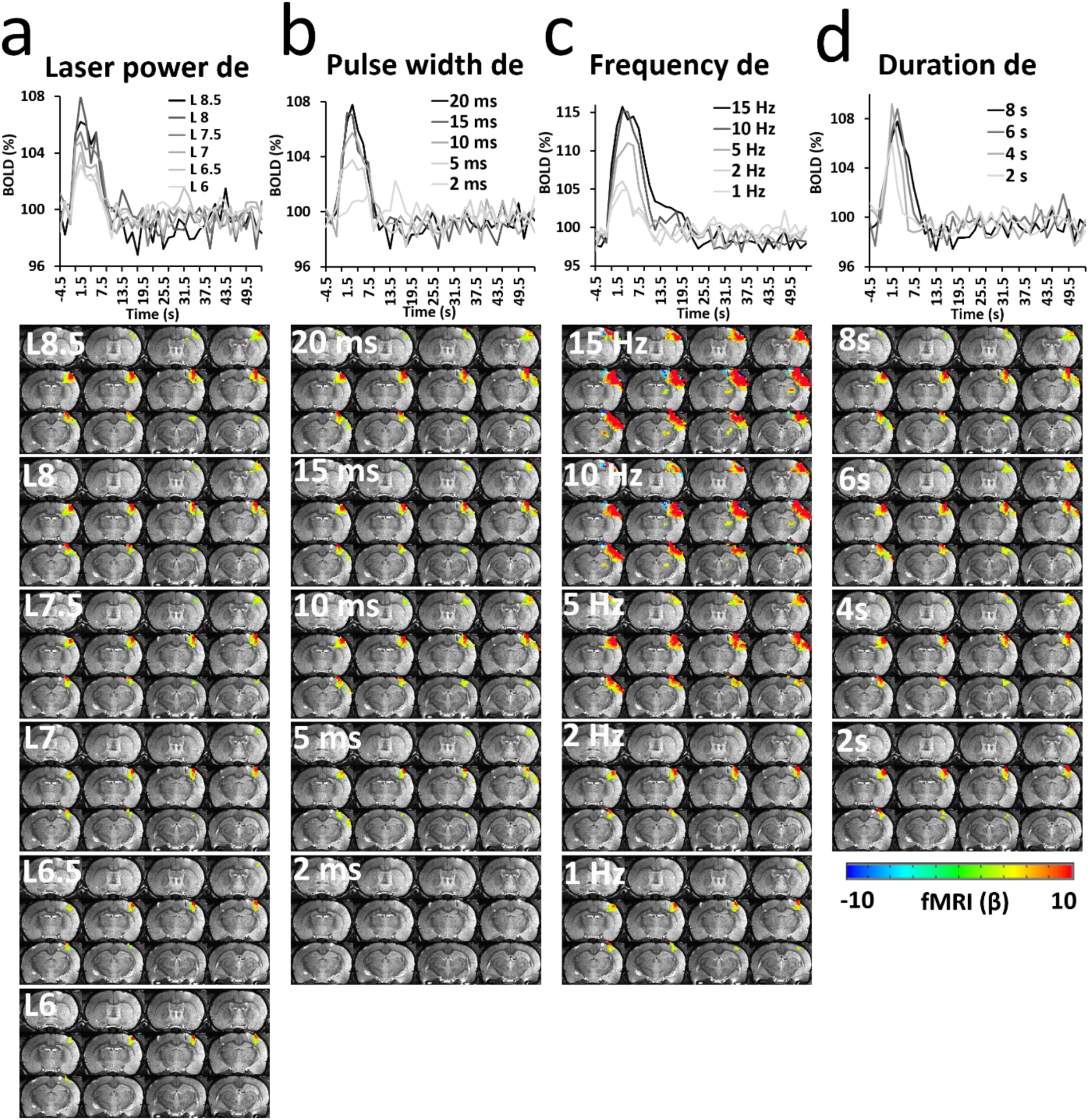
Representative functional maps and time courses of the fMRI signal (average of 13 epochs, 60 s per epoch) upon CC light activation with (**a**) laser power dependency (2 Hz, 8 s, 20 ms pulse width), (**b**) pulse width dependency (2, 5, 10, 15 and 20 ms pulse width, 2 Hz, 8 s, L 8), (**c**) frequency dependency (1, 2, 5, 10 and 15 Hz, 8 s, L 8, 20 ms pulse width) and (**d**) duration dependency (2, 4, 6 and 8 s, 2 Hz, 20 ms pulse width, L 8). It is noteworthy that exposure to light with high frequency (10 and 15 Hz) at high power (35 mW) led to heating effects, inducing artifacts close to the fiber tip (**c**), as well as very strong antidromic activity. GLM-based t-statistics in AFNI is used, p (corrected) < 0.005.

**Supplementary Figure 2.**
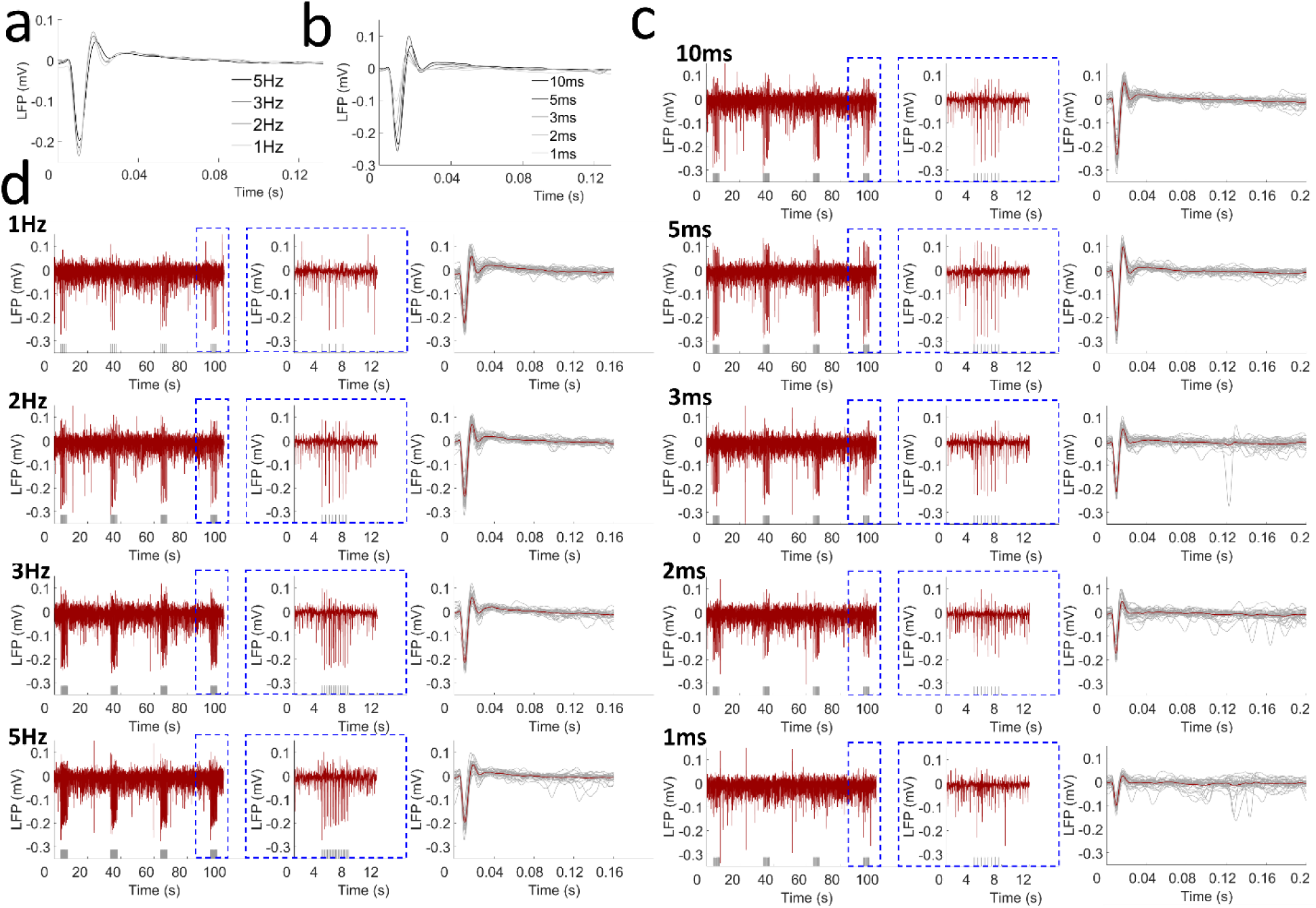
The light-driven antidromic LFP with frequency and pulse width dependency of a representative rat. (**a**) Averaged LFP driven by light pulses at different frequencies (1, 2, 3 and 5 Hz; 10 ms pulse width, L 7.5, 4 s stimulation 26 s rest, 16 epochs). (**b**) Averaged LFP driven by light pulses at different pulse widths (1, 2, 3, 5 and 10 ms pulse width; 2 Hz, L 7.5, 4 s stimulation 26 s rest, 16 epochs). (**c**) The raw LFP trace by optogenetic stimulation (*left*, 4 epochs), the enlarged representative LFP for one epoch (*middle*) and the averaged LFP from one trial (red line).The grey lines show all the LFP from this trial (*right*) upon different stimulation frequencies. (**d**) The raw LFP trace during optogenetic stimulation (*left*, 4 epochs), the enlarged representative LFP for one epoch (*middle*) and the averaged LFP from one trial (red line). The grey lines show all the LFP from this trial (*right*) upon different stimulation light pulse widths. Grey lines beneath the LFP indicate the stimulation.

**Supplementary Figure 3.**
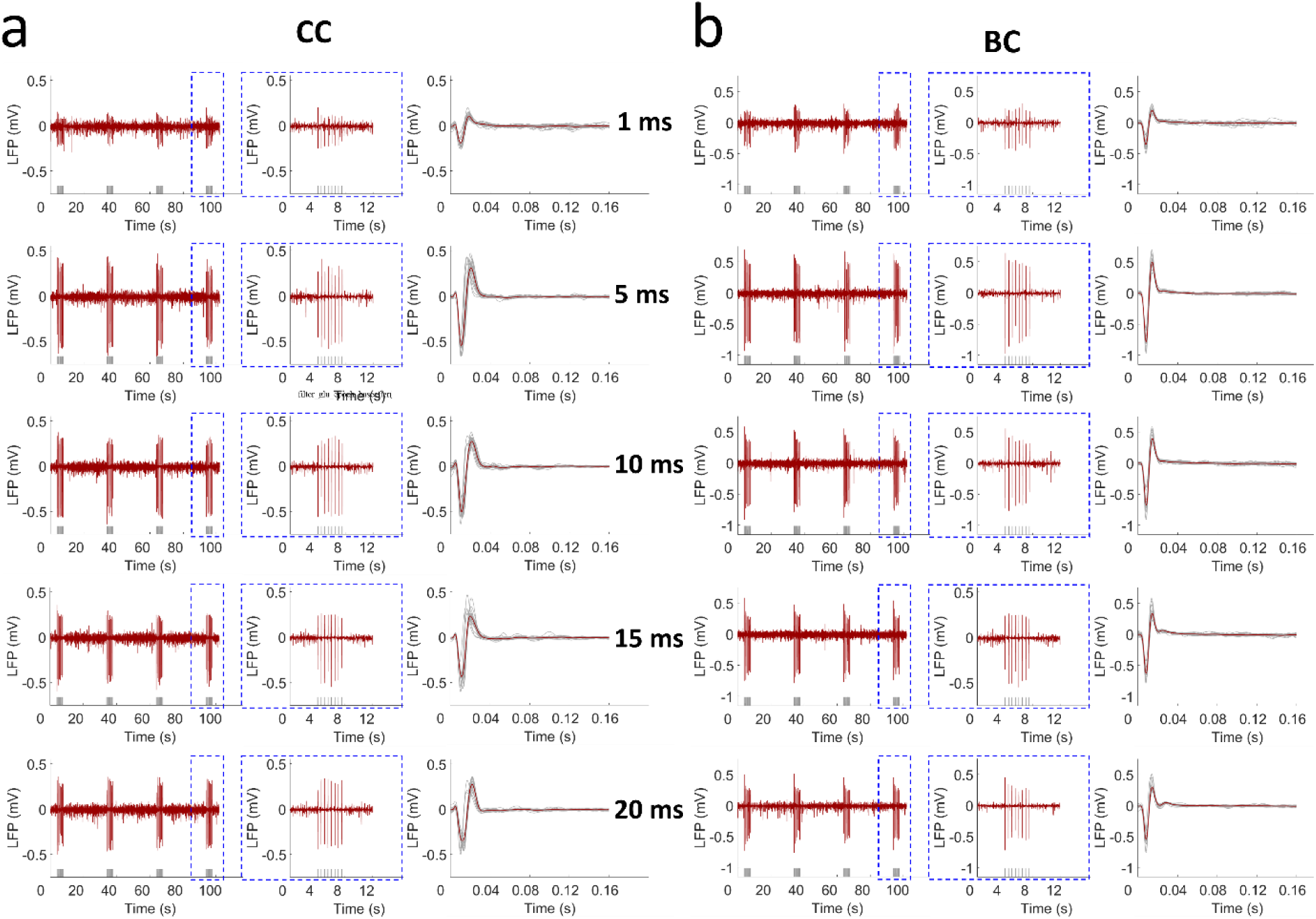
Light-driven LFP for antidromic activity from CC (**a**) stimulation and BC (**b**) direct stimulation showing similiar pattern with pulse width dependency of a representative rat. Every panel in **a** and **b** shows the raw LFP trace observed upon optogenetic stimulation (*left*, 4 epochs), the enlarged representative LFP for one epoch (*middle*) from the dashed blue box and the averaged LFP from one trial (red line). The grey lines show all the LFP from this trial (*right*). Grey lines beneath the LFP indicate the stimulation.

**Supplementary Figure 4.**
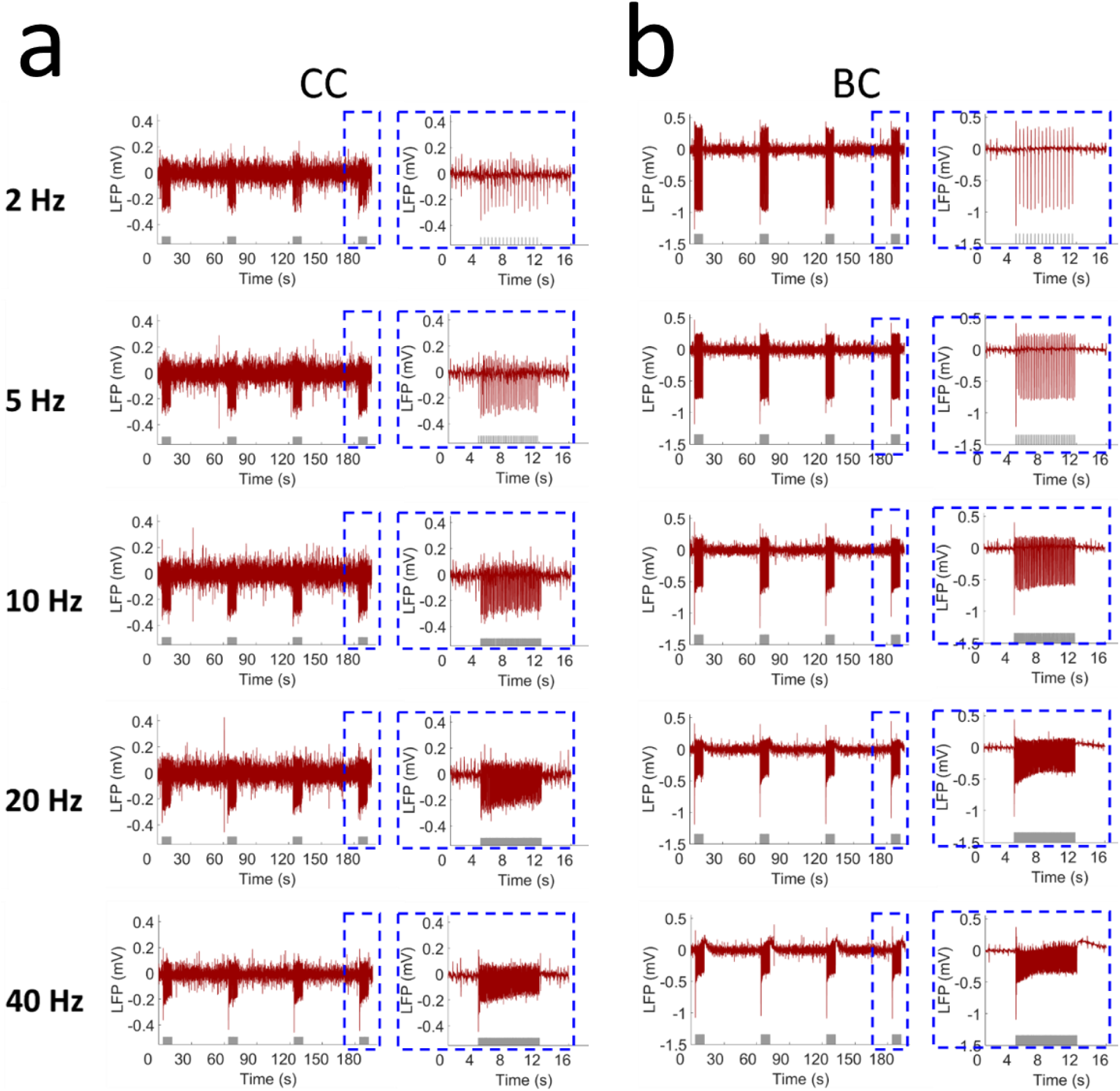
Light-driven LFP for antidromic activity from CC stimulation (**a**) and BC direct stimulation (**b**) showing similar pattern with frequency dependency of a representative rat. Every panel in **a** and **b** shows the raw LFP trace by optogenetic stimulation (*left*, 4 epochs), the enlarged representative LFP for one epoch (*right*) from the dashed blue box. Grey lines beneath the LFP indicate the stimulation.

**Supplementary Figure 5.**
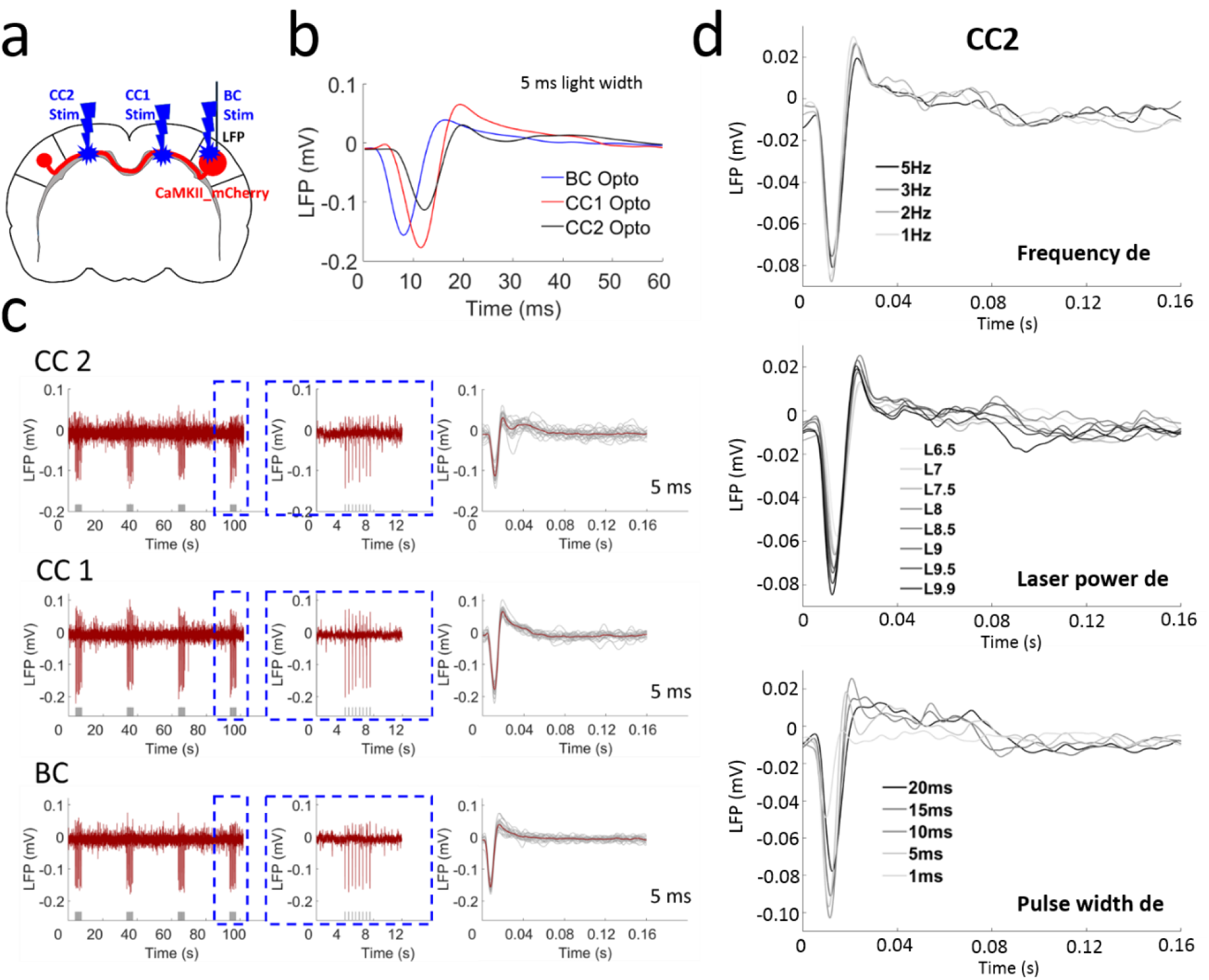
Light-driven LFP for antidromic activity from CC stimulation in both hemispheres and BC direct stimulation. (**a**) The schematic plan for the experiment design. (**b**) Averaged LFP from the CC2 stimulation in the hemisphere opposite to the virus injection site (blue line), CC1 stimulation in the same hemisphere (red line) and BC direct stimulation (black line) shown different temporal features. (**c**) The raw LFP trace by optogenetic stimulation (*left*, 4 epochs), the enlarged representative LFP for one epoch (*middle*) from the dashed blue box and the averaged LFP from one trial (red line). The grey lines show all the LFP from this trial (*right*). (**d**) Averaged LFP upon optogenetic stimulation of CC2 with frequency (*upper panel*), laser power (*middle panel*) and pulse width (*lower panel*) dependency showing reliably detected antidromic activity.

**Supplementary Figure 6.**
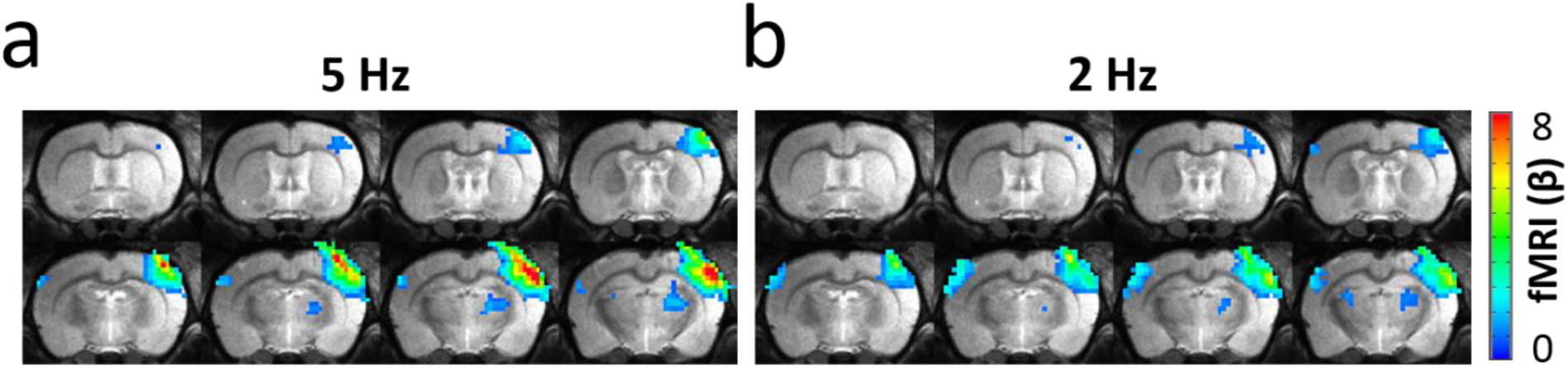
Light-driven functional maps demonstrating opposite relationships for antidromic and orthodromic activities in the BC to 5 Hz (a) and 2 Hz (b). The antidromic activity in the right hemisphere and the orthodromic activity in the left hemisphere responses to 5 Hz was stronger and weaker, respectively, compared to 2 Hz. GLM-based t-statistics in AFNI is used. p (corrected) < 0.01.

**Supplementary Figure 7.**
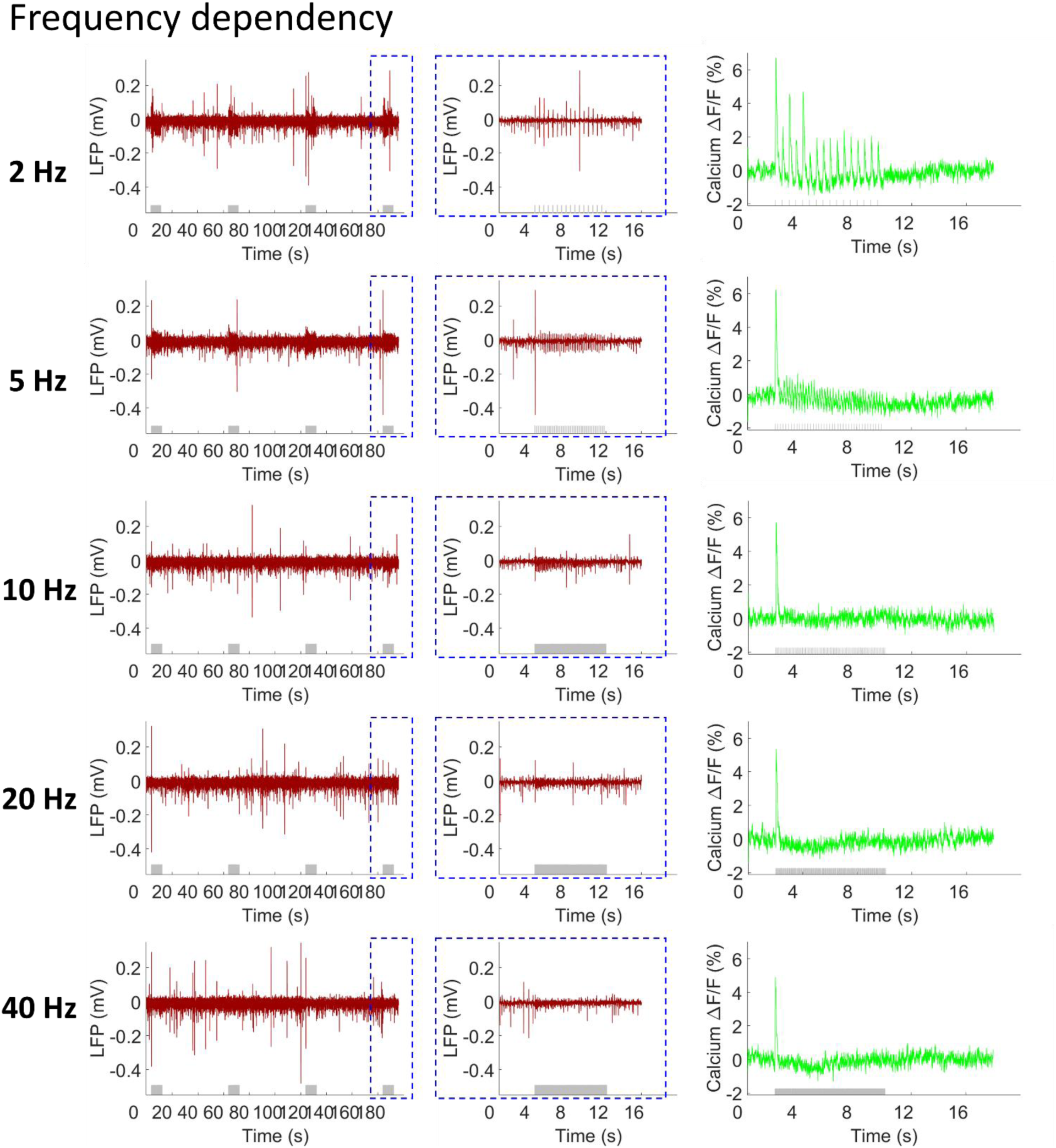
The frequency dependency of simultaneous LFP (red) and calcium response signals (green). Every panel shows the raw LFP trace elicited by optogenetic stimulation (*left*, 4 epochs), the enlarged representative LFP for one epoch from the dashed blue box (*middle*) and averaged calcium signal (8 s stimulation 52 s rest, 15 epochs, L9, pulse width 10 ms). Grey lines beneath the LFP and calcium signals indicate the stimulation.

**Supplementary Figure 8.**
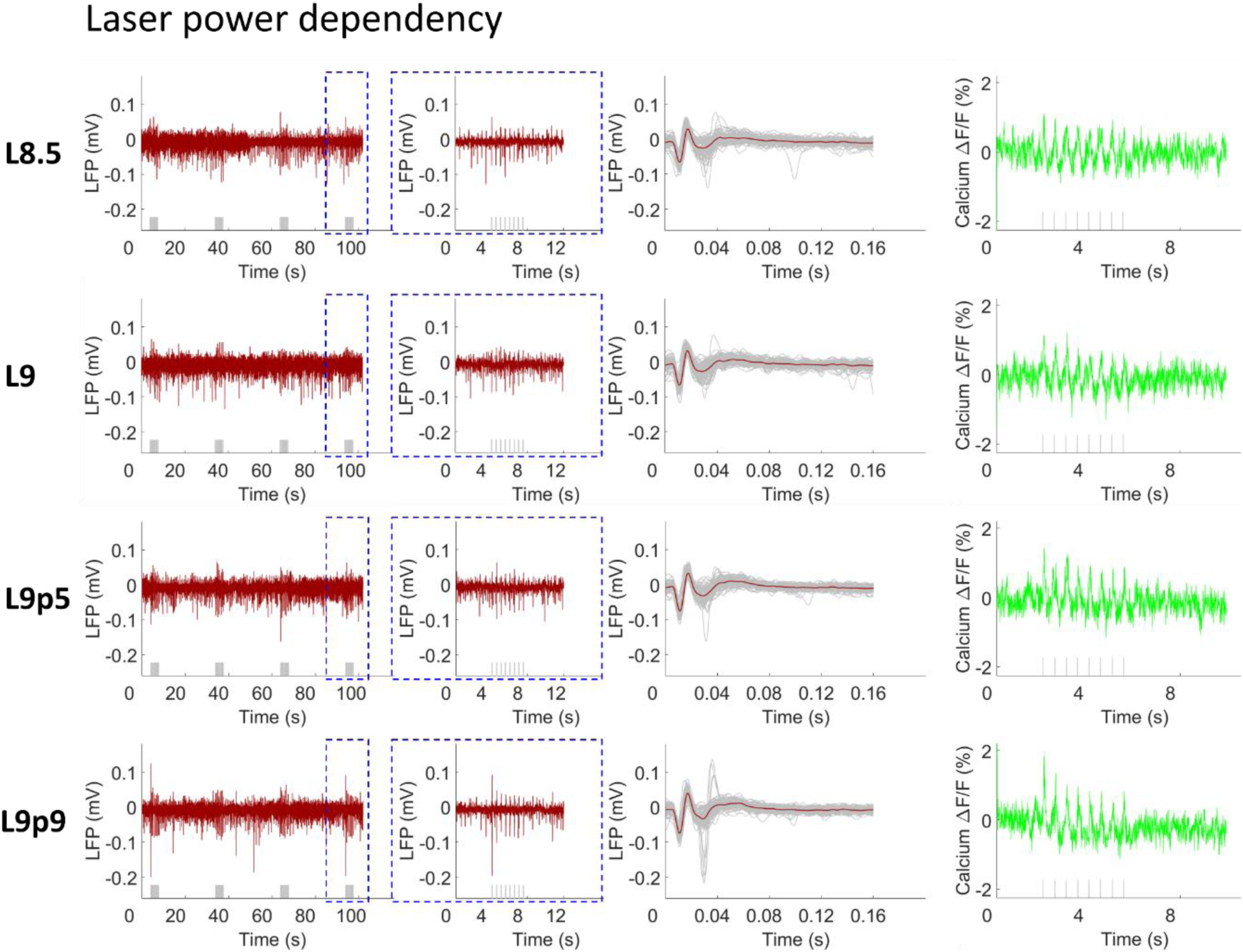
The laser power dependency of simultaneous LFP (red) and calcium signals (green) showing that both amplitudes increased as a function of the laser power. Every panel shows the raw LFP trace elicited by optogenetic stimulation (*left*, 4 epochs), the enlarged representative LFP for one epoch from the dashed blue box (*middle*), the averaged LFP from one trial (red line). Grey lines showing all the LFP from this trial (*right*) and the averaged calcium signal (4 s stimulation 26 s rest, 11 epochs, L9, pulse width 10 ms). Grey lines beneath the LFP and calcium signals indicate the stimulation.

**Supplementary Figure 9.**
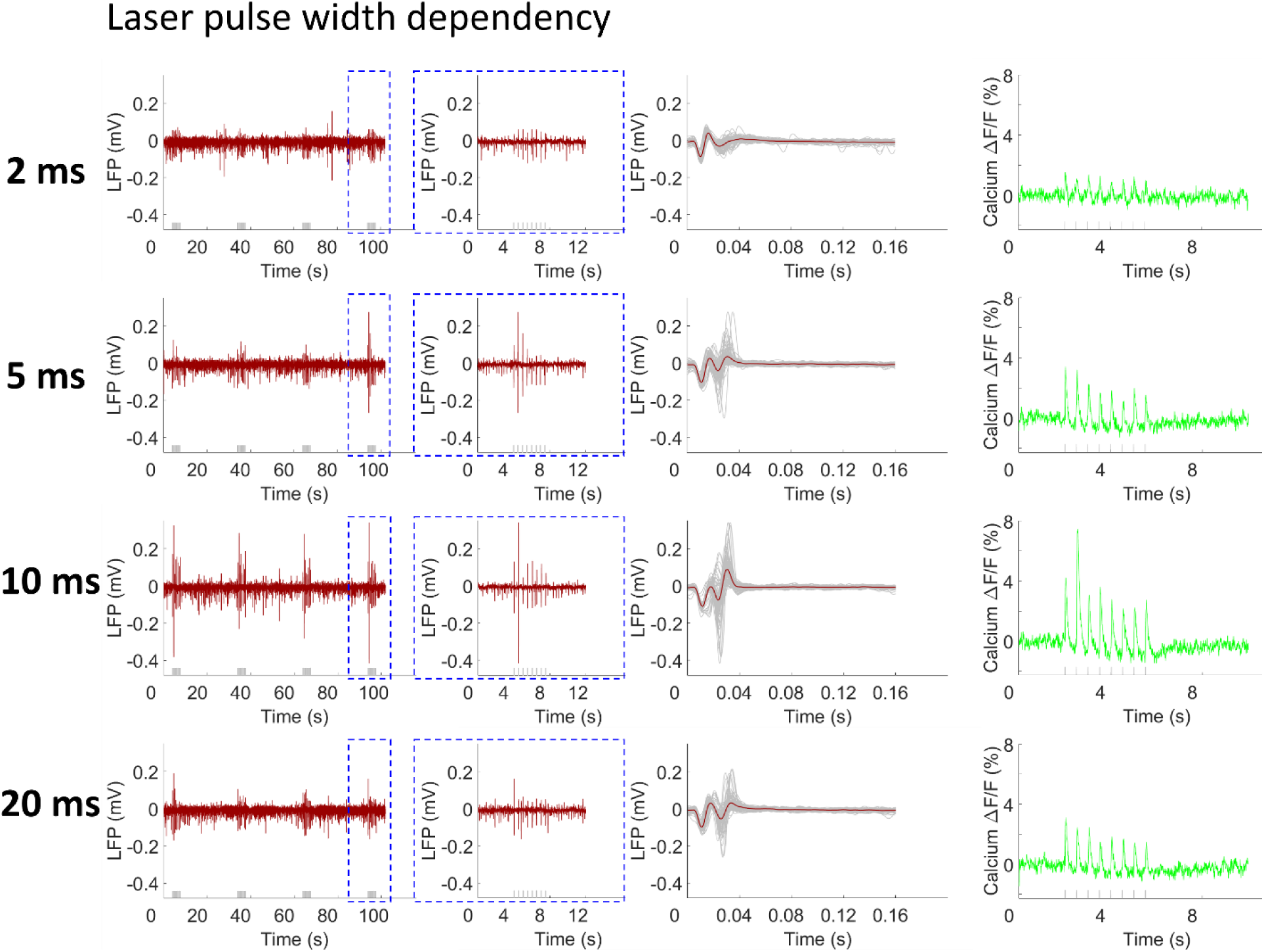
The laser pulse width dependency of simultaneous LFP (red) and calcium signals (green). Calcium signals increased and the LFP pattern demonstrated stronger depolarization according to the increased pulse width at 2 ms, 5 ms and 10 ms. In contrast, for the 20 ms pulse width stimulation, there was decreased calcium signal and weaker depolarization of LFP. Every panel shows the raw LFP trace elicited by optogenetic stimulation (*left*, 4 epochs), the enlarged representative LFP for one epoch from the dashed blue box (*middle*), the averaged LFP from one trial (red line, with the grey lines showing all the LFP from this trial) (*right*) and the averaged calcium signal (2 Hz, 4 s stimulation 26 s rest, 11 epochs, L9). Grey lines beneath the LFP and calcium signals indicate the stimulation.

**Supplementary Figure 10.**
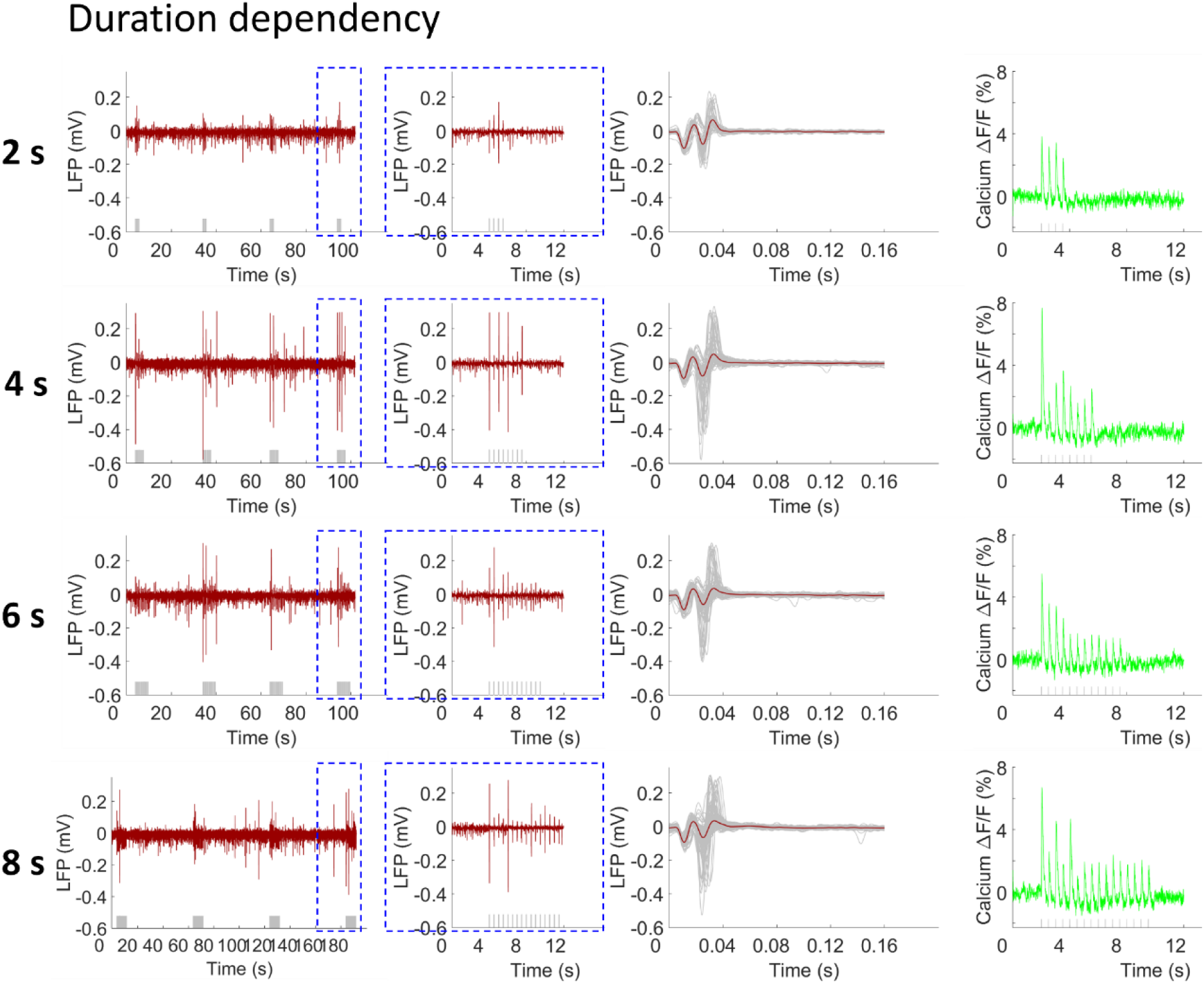
The duration dependency of simultaneous LFP (red) and calcium signals (green). Every panel shows the raw LFP trace elicited by optogenetic stimulation (*left*, 4 epochs), the enlarged representative LFP for one epoch from the dashed blue box (*middle*), the averaged LFP from one trial (red line, with the grey lines showing all the LFP from this trial) (*right*) and the averaged calcium signal (2 Hz, 11 epochs, L9, pulse width 10 ms). Grey lines beneath the LFP and calcium signals indicate the stimulation.

**Supplementary Figure 11.**
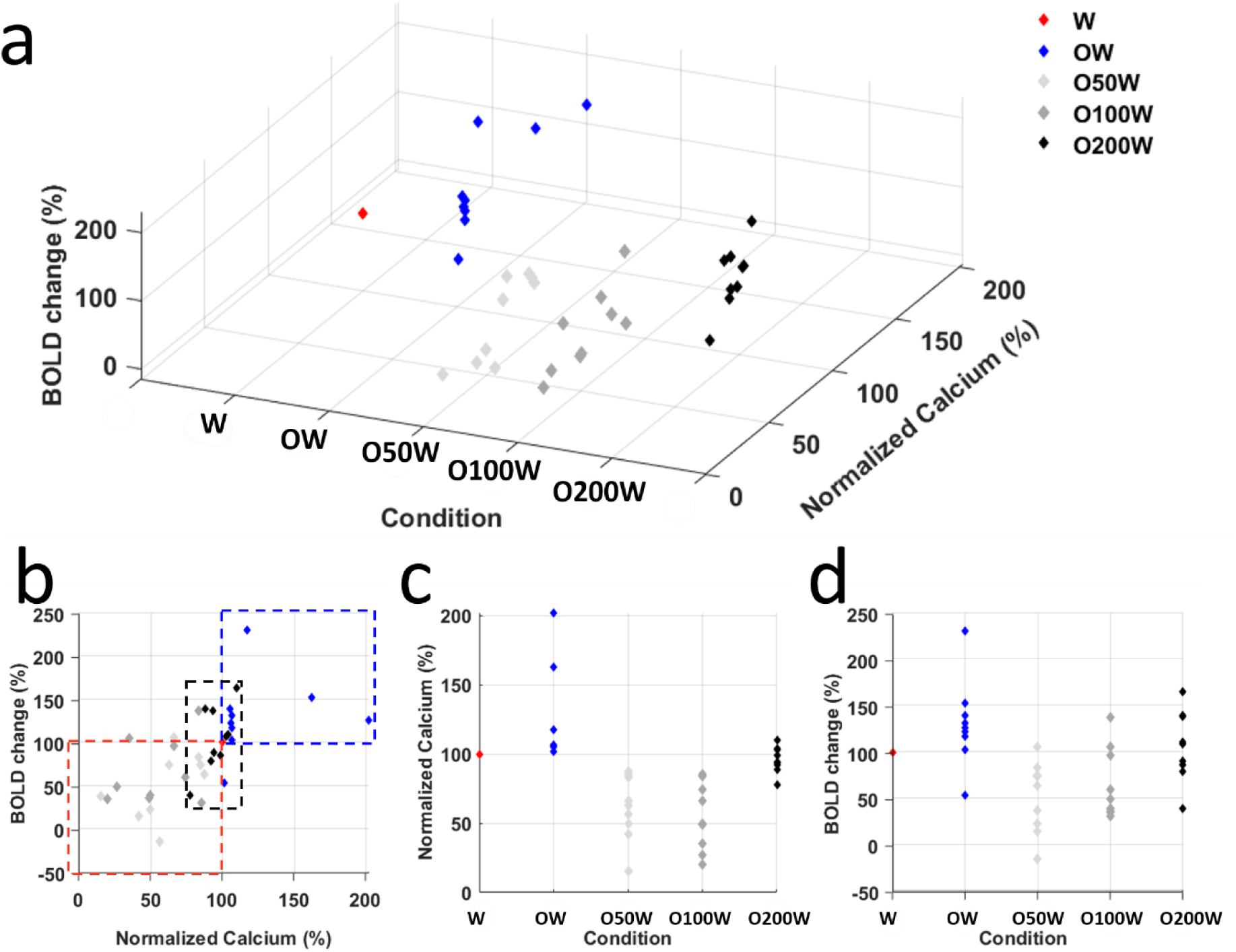
The scatter plots of the evoked BOLD and calcium signals for 5 stimulation conditions (W, OW, O50W, O100W and O200W) in 9 animals. (**a**) 3D plot of the BOLD changes (Z axis), calcium changes (Y axis) and stimulation conditions (X axis). Both BOLD and calcium signals are normalized to condition W. (**b**) View from the correlation of BOLD changes with calcium signals. The central red diamond is the baseline to which the data were normalized. Blue diamonds represent the condition OW, most of them distributed in the dashed blue box, showing increased neuronal activities. Light grey diamonds and dark grey diamonds represent the condition O50W and O100W, respectively, most of them located in the dashed red box, showing suppressed neuronal activities. (**c**) Normalized calcium signals as a function of condition. (**d**) Normalized BOLD changes as a function of condition. W: whisker stimulation only, OW: simultaneous optical and whisker stimulation, O[x]W optical stimulation followed by [x] ms-delayed whisker stimulation.

**Supplementary Figure 12.**
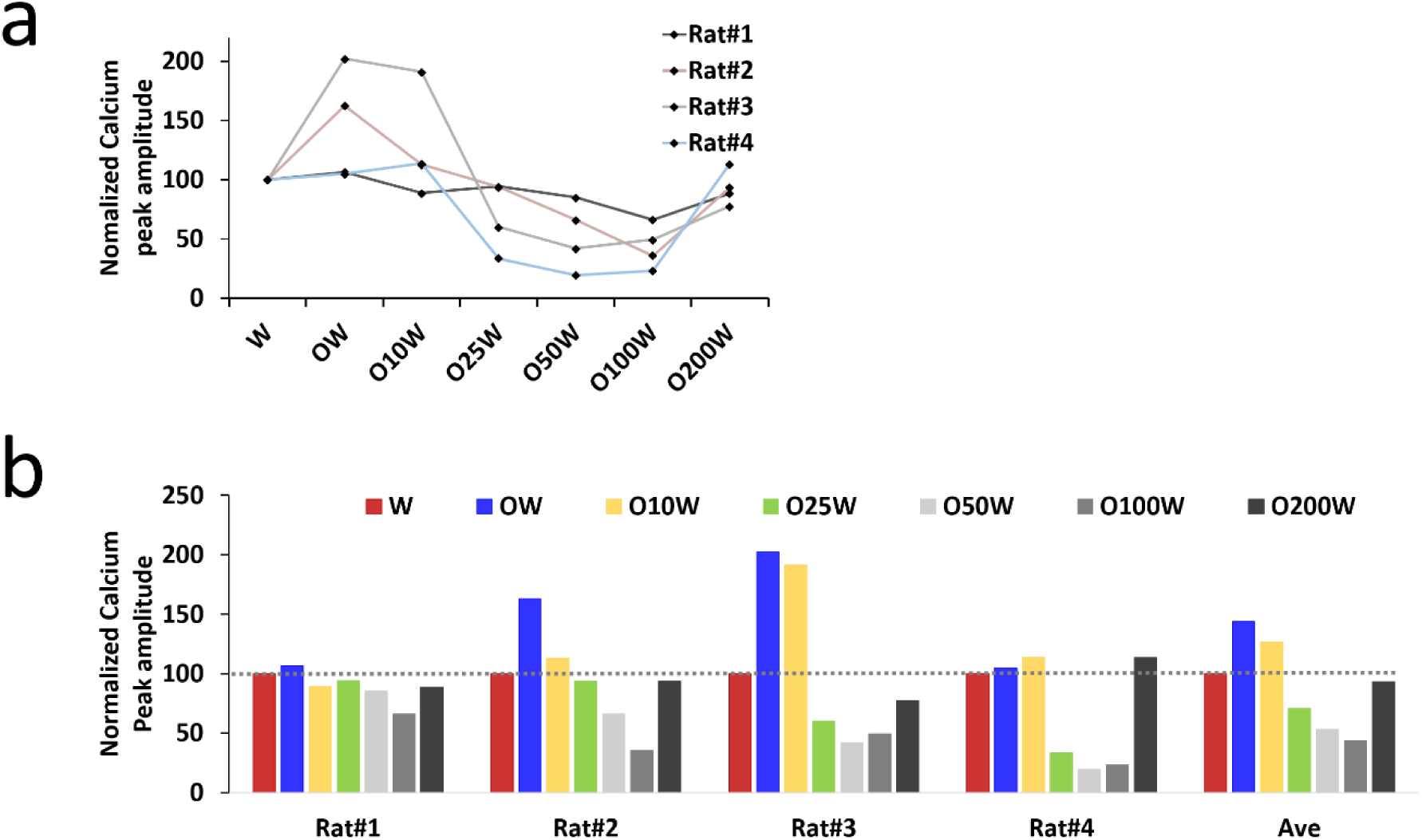
The effect of conditioning stimuli in the sensory evoked calcium signals in the left hemisphere for 7 refined conditions (W, OW, O10W, O25W, O50W, O100W and O200W). (**a**) The scatter plot of the calcium signals normalized to condition W from 4 rats. (**b**) The individual pattern changes of calcium signals from 4 animals, as well as the averaged calcium signal change pattern for all the 7 conditions.

**Supplementary Figure 13.**
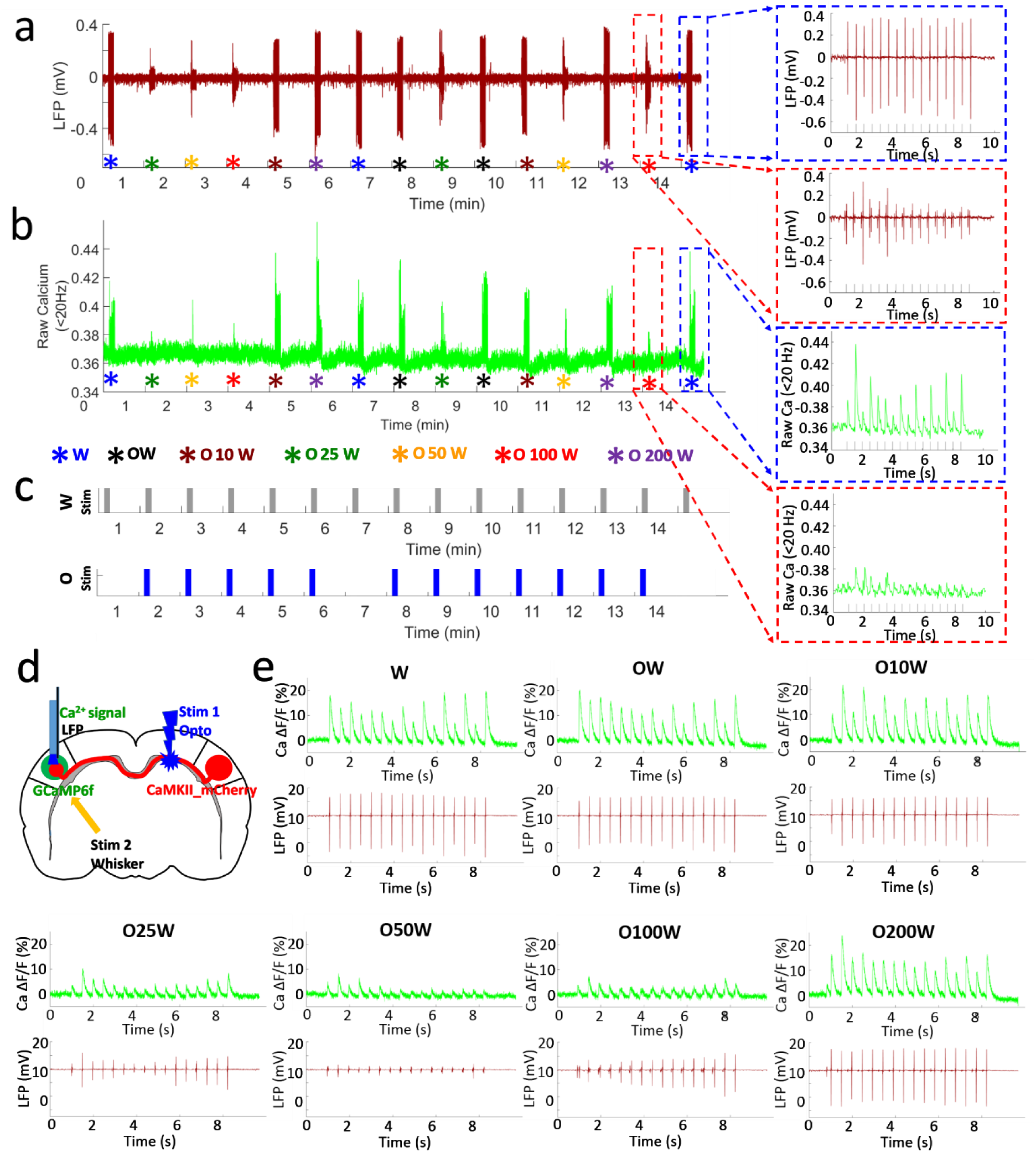
Typical LFP (red) and calcium signals (green) of one trial from a representative rat. (**a**) Different amplitudes of LFP and (**b**) calcium signal changes showing the different neuronal activity upon seven randomized stimulation conditions. (**c**). Simplified diagram representing the typical calcium signals and LFP for condition W (blue dash boxes in **a** and **b**, upper graph in **c**) and O100W (red dash boxes in **a** and **b**, lower graph in **c**) in one epoch. (**d**) Schematic of the experimental design. (**e**) Averaged calcium signals and LFP in left barrel cortex, further confirming the spatial and temporal features of sensory-evoked cortical activity pattern shaped by callosal inputs. W: whisker stimulation only, OW: simultaneous optical and whisker stimulation, O[x]W optical stimulation followed by [x] ms-delayed whisker stimulation.

**Supplementary Figure 14.**
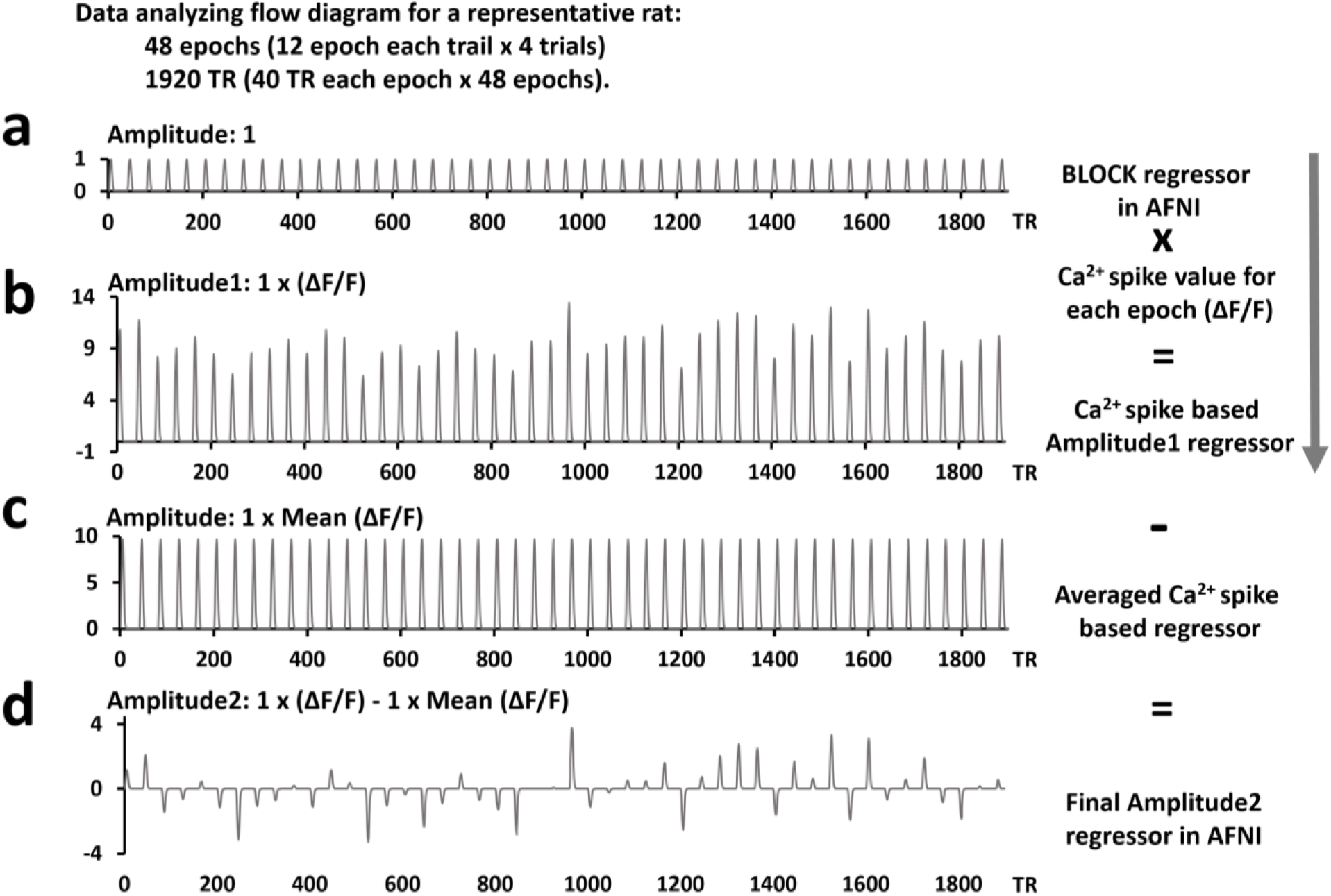
The flow diagram to generate the calcium signal-based regressor for the fMRI correlation map. (**a**), version 1 of the regressor, generated with the parameter BLOCK (L, 1), which generates a convolution of a square wave of duration L with the stimulation train and makes a peak amplitude of block response = 1. (**b**), the variable calcium amplitude of each epoch from a representative rat is used to generate the AM1 (amplitude modulated 1) regressor in 3dDeconvolve command in AFNI. (**c**), the averaged calcium amplitude of all the epochs is used to generate the regressor of no interest. (**d**), by computing ‘b – c’, the differences from the mean calcium amplitude can be detected. This new vector constitutes the final regressorAM2. ‘AM2’ allows to detect voxels that activate but do not change proportionally to the amplitude factor, as well as provides a direct measure of the proportionality of the activation in response to changes in the input amplitude factors (from the description of 3dDeconvolve program in AFNI).

